# A ratio-based framework using Quartet reference materials for integrating long- and short-read RNA-seq

**DOI:** 10.1101/2025.09.15.676287

**Authors:** Qingwang Chen, Xiaorou Guo, Duo Wang, Jiaxin Zhao, Yang Xu, Yupei You, Yuanbang Mai, Shumeng Duan, Yaqing Liu, Yutong Zhang, Xiaojing Li, Hu Chen, Wanwan Hou, Ying Yu, Lianhua Dong, Jinming Li, Matthew E. Ritchie, Rui Zhang, Leming Shi, Yuanting Zheng

## Abstract

Long-read RNA sequencing (lrRNA-seq) enables full-length transcript profiling but is confounded by technical batch effects that compromise quantification and prevent data integration across platforms, protocols, and laboratories. The lack of a transcriptome-wide biological ground truth has hindered objective benchmarking. To address these dual challenges, we leveraged certified Quartet reference materials to generate one of the largest multi-center lrRNA-seq resources to date: over one billion long reads from 144 libraries across four PacBio and Nanopore protocols in four independent laboratories. We first establish that ratio-based quantification against built-in reference samples effectively removes technical noise, revealing underlying biological signals. We then constructed the first ratio-based reference datasets for full-length transcripts— comprising 10,218 isoforms and 6,032 alternative splicing (AS) events—and orthogonally validated them with RT–qPCR. Finally, a comprehensive benchmark using these ground truths reveals that a hybrid strategy integrating long- and short-read data (hybrid-seq) achieves the highest quantification accuracy for both isoforms and AS events. Our work provides a foundational framework and resource for evaluating lrRNA-seq technologies and accelerating the standardization of full-length transcriptomics for research and clinical applications.

---

**Isoforms** are distinct mRNA transcripts derived from a single gene through mechanisms such as alternative splicing (AS), which significantly expands proteomic diversity and regulates cellular function^1^. Aberrant splicing is a key disease mechanism, generating pathogenic isoforms that drive tumour growth and metastasis^2^, while germline splicing defects underlie cardiomyopathies, muscular dystrophies, and other hereditary diseases^3^. Each isoform is defined by a unique splice junction chain—a specific series of exon-exon junctions^4^. AS represents differential use of these junctions, exons, or splice sites, quantified by shifts in junction usage (e.g., percent spliced in, PSI) and the consequent redistribution of isoform abundances^5^. Critically, because junction usage constrains isoform identification and isoform abundances aggregate to gene-level expression^6^, accurate profiling across all three interconnected layers—gene, isoform, and junction—becomes progressively more challenging at finer resolutions. This multi-level accuracy is essential for deriving mechanistic insight, discovering biomarkers, and enabling precision therapeutics^7^.

The advent of long-read RNA-seq (**lrRNA-seq**) technologies, such as Pacific Biosciences (PacBio) HiFi and Oxford Nanopore Technologies (ONT) duplex sequencing, has ushered in an era of full-length transcriptomics. A key advantage of lrRNA-seq is its ability to span entire splice junctions within a single read. This capability significantly improves the detection of AS events and the identification of full-length isoforms, while largely eliminating the need for computational reconstruction and reducing ambiguity in assigning shared exons^8^. In contrast, short-read RNA-seq (srRNA-seq) fragments transcripts into 50–300 nt pieces, requiring complex computational assembly to infer isoform structures from junction-spanning read pairs^9^. The “one-read-one-transcript” nature of lrRNA-seq typically yields more unambiguous isoform architectures and abundances, particularly for transcripts with multiple junctions or complex splicing patterns^10^. Furthermore, recent advances in sequencing chemistry and base-calling algorithms have elevated lrRNA-seq per-base accuracy to near-short-read levels (PacBio HiFi ≈ Q30, ∼99.9%; ONT duplex > Q20, ∼99%)^7^, thereby coupling long reads with high fidelity to enhance both qualitative resolution and quantification accuracy.

However, the absolute quantification of isoforms remains highly susceptible to **batch effects** stemming from differences in platforms, protocols, laboratories, and analytical pipelines^11^. The root cause lies in the reliance on run-specific measures like read counts or counts per million (CPM), which are assumed to be directly comparable across experiments but instead accumulate technical variation at every step^12^. This includes disparities between platforms (PacBio vs. ONT), library protocols (ISO-Seq, MAS-ISO-Seq, cDNA, and direct-RNA), laboratory procedures, and bioinformatic tools (aligner and quantifier), resulting in inconsistent absolute counts across batches. These systematic variations mirror the batch effects previously documented for gene-level quantification in short-read RNA studies^13,14^. Existing lrRNA-seq benchmarks—such as LRGASP^15^, SG-NEx^10^, LongMix^16^, and various spike-in or simulation efforts^17,18^— have been invaluable for comparing protocols and analytical pipelines, revealing protocol-specific biases (e.g., coverage and 3′ bias), the impact of read quality, and substantial pipeline-dependent variability in isoform detection and quantification. However, these pioneering efforts were primarily conducted in single or tightly coordinated environments. Consequently, they could not resolve the critical issue of cross-laboratory batch effects, which is essential for real-world applications and multi-center data integration. More importantly, the lack of a transcriptome-wide biological ground truth precluded definitive assessment of accuracy across platforms and laboratories. The absence of a multi-laboratory, multi-platform, multi-protocol benchmark remains the fundamental bottleneck obstructing data integration and the reliable application of long-read sequencing in clinical and industrial settings.

**Ratio**-based quantification against built-in reference samples represents a paradigm shift from absolute quantification in multi-omics profiling, effectively removing technical variations arising from platforms, protocols, or analytical pipelines^12,13^. This approach mitigates batch effects while preserving biological signals by normalizing the absolute quantifications of study samples to those of concurrently profiled reference materials on a feature-by-feature basis, i.e., the sample-to-reference ratio (SRR)^12,13^. This principle was first established and verified by the Quartet Project for integrating gene-level and splicing-level expression across multi-batch srRNA-seq data^13^, enabling unprecedented multi-batch data integration. Crucially, this framework provides the foundation for establishing a key component missing in previous benchmarks: a transcriptome-wide, biologically defined transcriptomics ground truth^19^. However, the transformative potential of this ratio-based framework for long-read transcriptomics remains entirely untested.

Despite these advances, three critical **gaps** impede the broad adoption of lrRNA-seq for routine isoform and AS profiling. First, the effectiveness of ratio-based quantification, though proven for genes and srRNA-seq, remains unverified for isoform-level integration across multi-batch lrRNA-seq data. Second, a transcriptome-wide, biologically validated ground truth is still lacking, which is essential for rigorous and reproducible benchmarking^20,21^. Third, existing benchmarks are narrow in scope, confined to single-laboratory assessments. Consequently, the latest lrRNA-seq protocols and analytical pipelines—including the hybrid-seq strategies^22,23^ that combine short- and long-read data within a unified framework—remain inadequately evaluated, offering limited practical guidance for the field. Establishing ratio-based reference datasets (ground truths) and conducting comprehensive multi-center benchmarking of state-of-the-art lrRNA-seq workflows are therefore essential to enable reliable, large-scale isoform and AS analysis.

To address these limitations, we leveraged certified Quartet RNA reference materials^12,13^ to generate one of the largest multi-center lrRNA-seq resources to date: over one billion long reads from 144 libraries. We demonstrate that ratio-based quantification successfully enables cross-batch integration of isoform expression data. Using a consensus framework applied to high-quality data, we constructed the first ratio-based reference datasets for isoforms and AS events, orthogonally validating them with RT-qPCR^24^. Together, the Quartet RNA reference materials and their accompanying reference datasets establish transcriptome-wide biological ground truths for benchmarking lrRNA-seq laboratories, platforms, protocols, and analytical pipelines, thereby accelerating standardization and promoting the broader adoption of long-read transcriptomics.

## Results Study design

To establish a comprehensive benchmark for lrRNA-seq, we generated large-scale, multi-laboratory, multi-platform, and multi-protocol datasets using certified Quartet RNA reference materials. These materials are derived from a genetically characterized Chinese quartet family, comprising monozygotic twin daughters (D5 and D6), their father (F7), and mother (M8)^12,13^.

We sequenced the Quartet RNA samples on both long-read (LR: PAB-PacBio and ONT-Oxford Nanopore Technologies) and short-read (SR: ILM-Illumina, MGI-Beijing Genomics Institute, and ELE-Element Biosciences) platforms^13,25^, generating 12 batches of lrRNA-seq and 22 batches of srRNA-seq data (**Fig. 1a**). The lrRNA-seq data were generated across four independent laboratories using four library preparation protocols: PAB MAS-ISO-Seq (Kinnex), PAB ISO-Seq, ONT PCR-cDNA, and ONT direct RNA (dRNA) (**Supplementary Table 1**). To enable precise assessment of pipeline performance, we spiked the Spike-In RNA Variant Set 4 (SIRV-SET4) into one MAS-ISO-Seq batch (**Supplementary Figs. 1-3**). Most libraries were designed to yield ≥5 million reads, a threshold sufficient for reliable isoform quantification^1,26^, resulting in one of the largest lrRNA-seq resources: over one billion long reads from 144 lrRNA-seq libraries across 12 batches (**Supplementary Fig. 4, Supplementary Table 2**). We also integrated 264 srRNA-seq libraries from nine laboratories^13,25^, comprising libraries prepared by poly(A) selection or ribosomal RNA depletion (RiboZ) for cross-technology comparison (**Supplementary Table 3**).

**Fig. 1.**
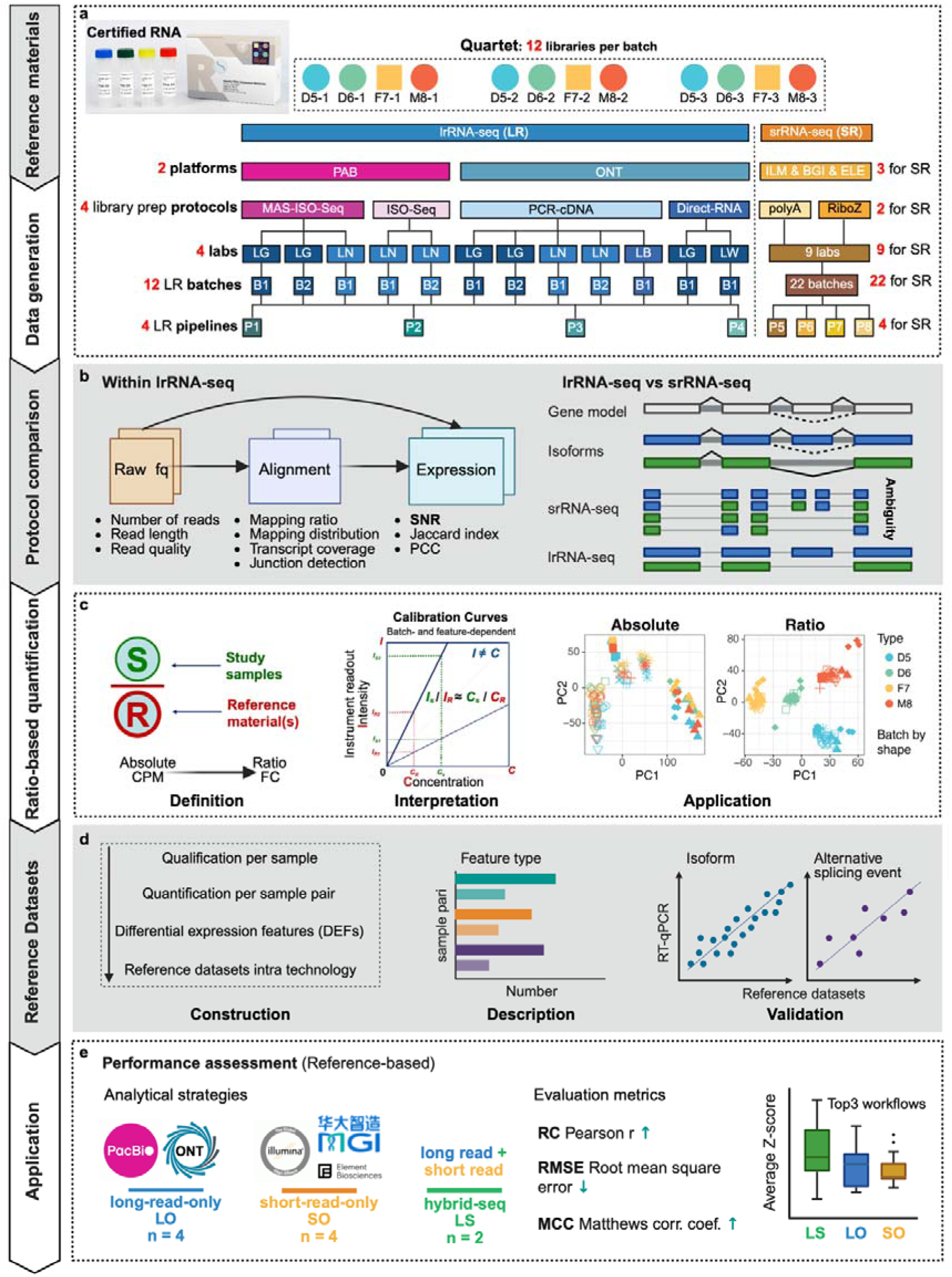
Overview of study design. **a**, Reference materials and data generation. Certified Quartet RNA reference materials were used for long-read (LR) RNA sequencing (lrRNA-seq) across multiple platforms, library preparation methods, and laboratories. These LR and short-read (SR) sequencing datasets were used for performance evaluation. **b**, Protocol evaluation. The lrRNA-seq data were assessed at three stages—raw reads, alignments, and expression—and contrasted with short-read data to quantify ambiguity of junctions and isoforms. Key metrics: read length/quality, mapping statistics, junction detection, signal-to-noise ratio (SNR), Jaccard index, and Pearson correlation coefficient (PCC). **c**, Ratio-based quantification. Absolute quantification of study samples (S) is scaled to concurrent reference materials (R) to yield ratio-based quantification (fold change, or sample-to-reference ratio, SRR). Ratios harmonize batches (PCA plots) and link instrument signal to concentration (calibration curve). **d**, Construction and validation of reference datasets. High-quality lrRNA-seq libraries were selected and integrated to construct ratio-based reference datasets for isoforms and AS events. Numbers of features per sample-pair are summarized; a subset was validated by RT–qPCR. **e**, Benchmark application. Performance of three analytical strategies—long-read-only (LO, n = 4), short-read-only (SO, n = 4), and hybrid-seq (LS, long + short, n = 2)— was scored with reference-based metrics. Boxplots show average Z-scores; hybrid-seq workflows rank highest.

Our analysis framework evaluated data at three key stages: raw reads, alignments, and expression quantifications. We also compared the performance of LR versus SR in isoform detection (**Fig. 1b**). We applied ratio-based quantification to integrate multi-batch lrRNA-seq isoform data, significantly improving cross-batch comparability. From high-quality lrRNA-seq datasets, we constructed high-confidence ratio-based reference datasets for isoform and AS events (**Fig. 1c**). We orthogonally validated these reference datasets using RT-qPCR assays (**Fig. 1d**), establishing them as ground truths for objective evaluation of sequencing protocols and analytical pipelines across LR, SR, and hybrid (LR+SR) RNA-seq workflows (**Fig. 1e**).

### Protocol-dependent differences in data quality

We comprehensively evaluated data quality across the 12 lrRNA-seq batches at the raw-read, alignment, and expression levels, observing pronounced protocol-dependent differences (**Supplementary Fig. 5**).

Raw-read quality varied significantly by platform and protocol. PacBio libraries consistently yielded longer reads and higher per-base quality (Q-score) than ONT libraries. While most protocols achieved the target of ≥ 5 million reads, PacBio ISO-Seq produced <1 million high-fidelity (HiFi) reads, and ONT direct RNA excelled in throughput, with each library exceeding 10 million reads. Despite lower yield, ISO-Seq reads were the longest and most accurate (mean length ≥ 2 kb, mean Q-score > 30). Overall, read length and quality declined in the order: ISO-Seq > MAS-ISO-Seq > ONT direct RNA > ONT PCR-cDNA (**Supplementary Fig. 6, Supplementary Table 4**).

Alignment and mapping characteristics also revealed key trade-offs. PacBio ISO-Seq datasets achieved near-perfect mapping ratios (> 99%), the highest among all protocols (**Supplementary Fig. 7**). While exonic reads dominated all datasets, the level of intronic signal—an indicator of nascent transcript capture and novel junction discovery—varied considerably. ISO-Seq and MAS-ISO-Seq contained up to 25% intronic reads, in contrast to ONT direct RNA, which had less than 10% (**Supplementary Fig. 8**). We observed a strong protocol-specific gradient in 5′→3′ coverage bias. This bias was minimal in ONT PCR-cDNA libraries (median ∼1.0) but became progressively more severe through direct RNA, ISO-Seq, and MAS-ISO-Seq (median <0.5), reflecting increasing 5′ loss in reverse-transcription-intensive protocols (**Supplementary Fig. 9**). Junction detection consistency was high for annotated junctions across platforms (0.76–0.79) but was substantially lower for novel junctions, highlighting the ongoing challenge of confident novel splice junction discovery, particularly for ISO-Seq (**Supplementary Fig. 10**).

Expression-level reproducibility systematically declined as transcriptional resolution increased. From gene- to isoform- to AS-level analysis, the median Jaccard index dropped from 0.88 to 0.65 to 0.47, the median Pearson correlation coefficients (PCC) fell from 0.99 to 0.98 to 0.78, and the median signal-to-noise ratio (SNR) decreased from 16 to 13 to 8 (**Supplementary Figs. 11-12**), underscoring the growing challenge of accurate detection and quantification at finer resolution.

We ranked the 12 batches of lrRNA-seq data by isoform-level SNR (**Supplementary Figs. 13-14**) and retained the seven highest-quality batches (SNR ≥ 10) for downstream analysis. In these seven batches, a clear transcript-length-dependent performance pattern emerged. Both platforms showed limited power to discriminate very short isoforms (< 400 nt; SNR ∼5). ONT libraries maintained high SNR (≥ 10) through the sub-kilobase range, whereas PacBio accuracy was lower for mid-length transcripts (400–600 nt; SNR ∼7) but increased steadily with length, surpassing ONT for isoforms longer than ∼1.5 kb (**Supplementary Fig. 15**).

### lrRNA-seq enables unambiguous isoform detection and quantification

LrRNA-seq captures complete transcript molecules, providing a direct advantage over srRNA-seq by spanning multiple splice junctions within a single read and eliminating the need for computational assembly^10^. Our analysis of seven high-quality lrRNA-seq and five high-quality srRNA-seq datasets (**Supplementary Figs. 16-17**) confirmed that lrRNA-seq provides superior coverage across multiple junctions, particularly for transcripts containing three or more splice junctions (**Fig. 2a**). Given that over half of all reference transcripts harbor three or more splice junctions (**Supplementary Fig. 18**), this capability is essential for accurately characterizing the transcriptome’s full complexity.

**Fig. 2.**
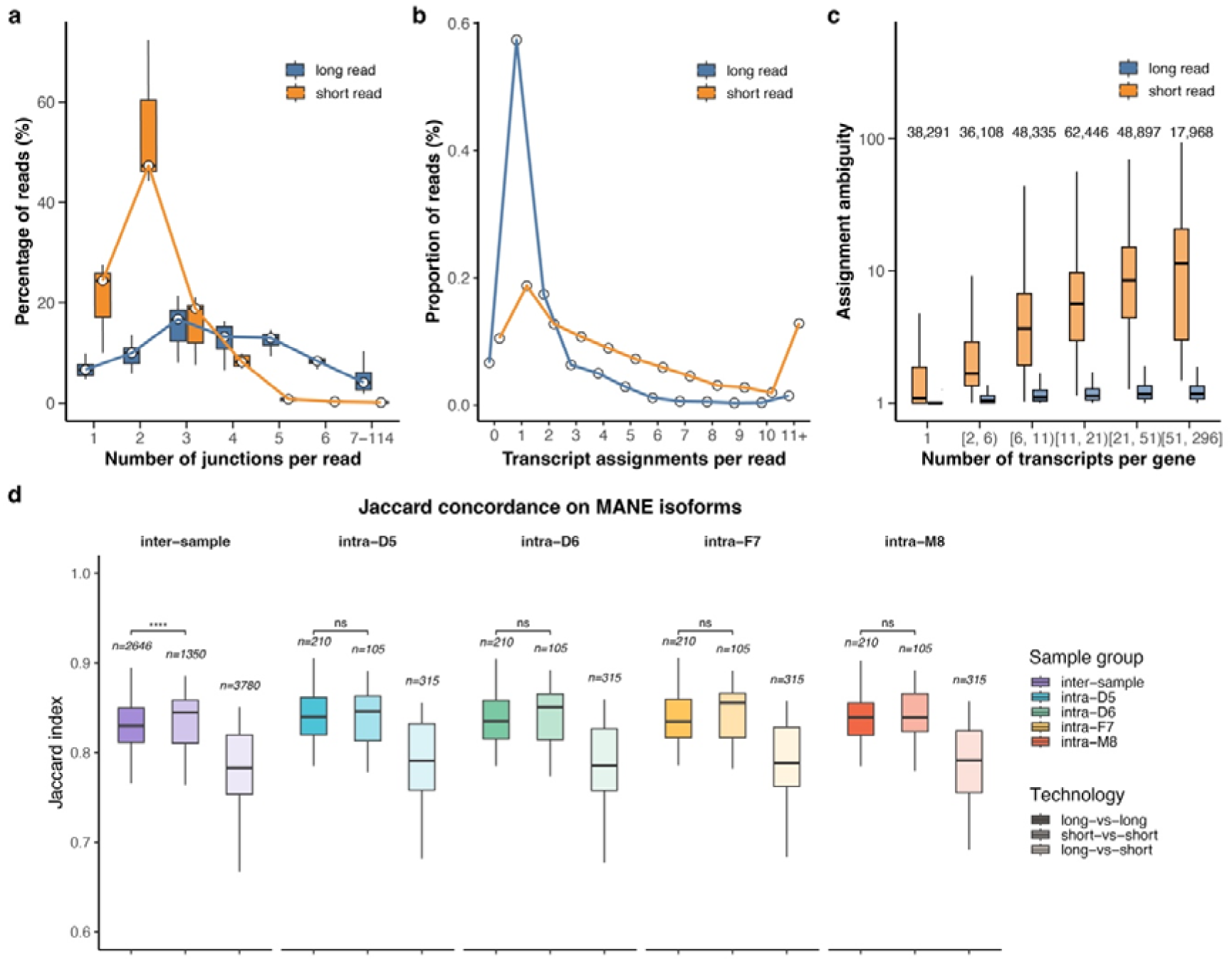
Long- versus short-read sequencing performance for isoform detection and quantification. **a**, Distribution of splice-junction coverage per read in long-read (blue) and short-read (orange) libraries; boxes show inter-quartile range (IQR) and whiskers 1.5 × IQR. **b**, Multiplicity of read-to-transcript assignments for the same datasets. **c**, Assignment ambiguity stratified by gene complexity (number of reference transcripts; values above boxes indicate isoform counts in each bin); y-axis on log scale. Values above each boxplot indicate the number of isoforms in that category. **d**, Jaccard index of intersect/union calls for MANE transcripts. Fill color encodes the sample group, and box-outline shade distinguishes the sequencing-technology pairing (dark = long vs. long, mid-grey = short vs short, light = long vs short). Boxes denote IQR; **** *P* < 1 × 10LL, ns = not significant (two-sided Wilcoxon rank-sum test).

A major advantage of lrRNA-seq is its substantial reduction of transcript assignment ambiguity^16^. We found that 64% of long reads mapped uniquely to a single transcript, in stark contrast to short reads, which frequently mapped ambiguously to many transcripts — often more than 11 per read (**Fig. 2b**). This lower mapping uncertainty was consistent across all multi-transcript genes, demonstrating the robustness of lrRNA-seq for accurate isoform detection and quantification (**Fig. 2c**).

We further evaluated isoform detection consistency using clinically curated Matched Annotation from NCBI and EMBL-EBI (MANE) isoforms^27^. The intra-sample detection of MANE isoforms was highly consistent. Remarkably, low-coverage lrRNA-seq achieved a median Jaccard index of 0.84, comparable to high-depth srRNA-seq. Inter-sample concordance was slightly higher for srRNA-seq (median = 0.86) than for lrRNA-seq (median = 0.83), likely due to its greater sequencing depth. However, concordance dropped significantly (median = 0.78) in direct cross-platform comparisons, indicating that technical differences between sequencing technologies— not biological variation—are the primary drivers of discordance in MANE isoform detection (**Fig. 2d**).

### Ratio-based isoform quantification enables data integration across batches

The integration of transcriptomic data from different sequencing technologies and batches remains a major challenge. Here, we demonstrate that ratio-based quantification is a powerful strategy for integrating isoform quantification data from lrRNA-seq and srRNA-seq. While absolute quantification showed high cross-platform consistency, technical differences overwhelmingly obscured inherent biological variation among sample types (**Supplementary Fig. 19a**). In contrast, ratio-based quantification effectively minimized this technical variability, resulting in substantially higher concordance among samples and revealing true biological signals (**Supplementary Figs. 19b-c**).

To quantify the extent of technical bias, we analyzed absolute quantification from 12 lrRNA-seq batches using the Oarfish pipeline^28^. Principal Component Analysis (PCA) analysis showed that samples clustered primarily by library protocols and sequencing platform, not by biological sample type (**Fig. 3a**). Principal Variance Component Analysis (PVCA) confirmed that these technical factors accounted for 72% of the total variance (**Fig. 3b**). Conversely, ratio-based isoform quantification dramatically reduced this technical noise, resulting in clear clustering by biological sample type (**Fig. 3c**) and establishing biological identity as the dominant source of variation (**Fig. 3d)**.

**Fig. 3.**
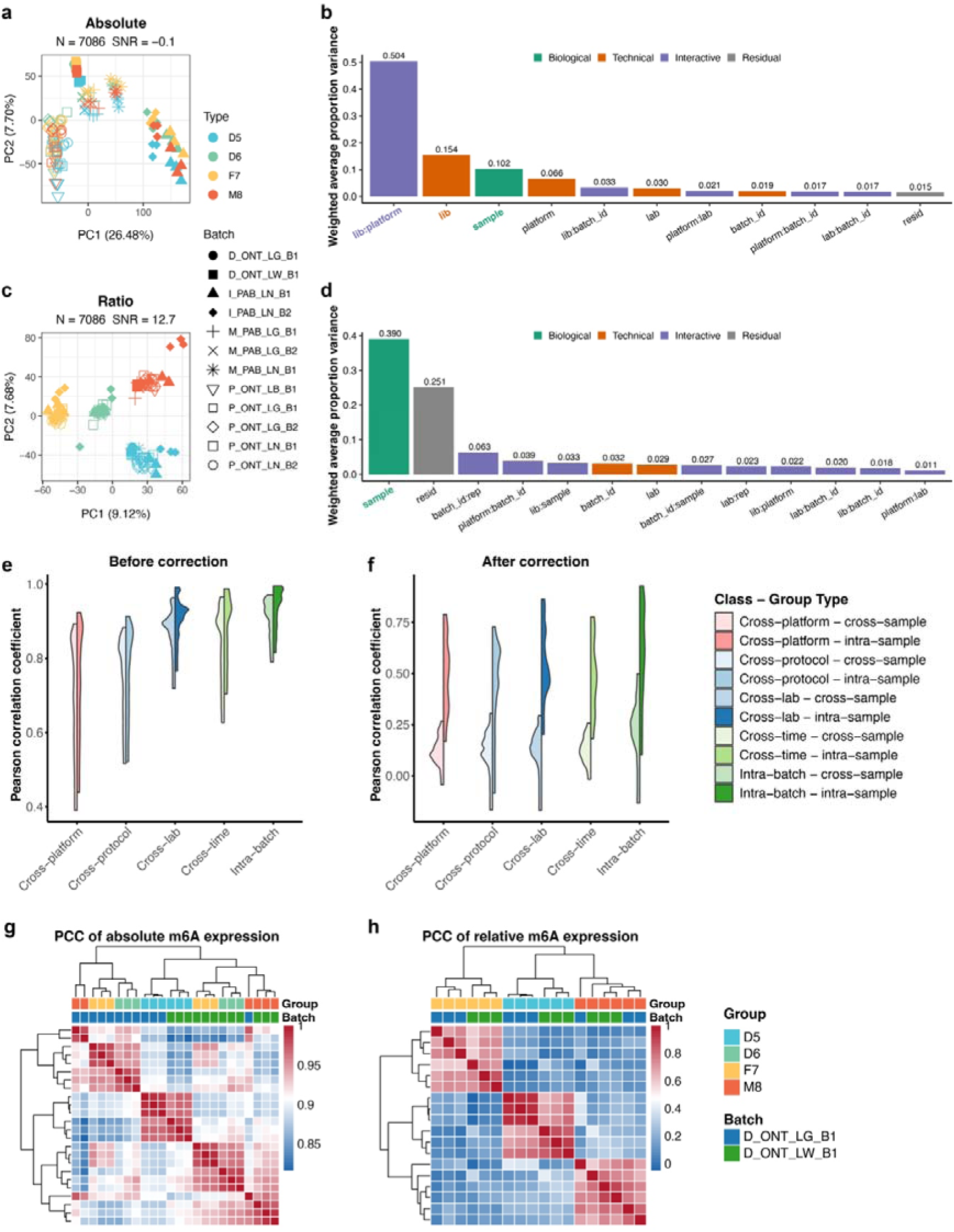
Ratio-based quantification improves the integration of isoform datasets. **a**, Principal Component Analysis (PCA) plot using absolute isoform quantification values. **b**, Principal Variance Component Analysis (PVCA) using absolute isoform quantification values. **c**, PCA plot using ratio-based isoform quantification values. **d**, PVCA using ratio-based isoform quantification values. **e**, Pearson correlation coefficients of isoform expression values across multiple conditions (cross-platform, cross-protocol, cross-lab, cross-time, and intra-batch) before ratio-based correction. **f,** Pearson correlation coefficients after ratio-based correction. **g**, Sample-to-sample PCC for absolute mLA quantification (site-level counts per million). Tiles are colored 0-1 (blue → red), annotation bars indicate sample (color code as in **a**) and batch (blue vs. green). PCC refers to Pearson correlation coefficient. **h**, PCC for ratio-based mLA quantification (within-batch ratio to the geometric mean of D6 samples). Ratio-based quantification removes most batch effects, causing biological groups to cluster together.

Importantly, this technical variability was not limited to sequencing platforms but also stemmed from bioinformatic pipelines. When integrating absolute isoform expression data from four analytical pipelines, the choice of pipeline itself became the dominant source of variance (**Supplementary Figs. 20a-b**). Applying ratio-based quantification removed this pipeline-specific bias and residual batch effects, enabling robust clustering by biological sample type and facilitating the construction of a transcriptome-wide reference dataset (**Supplementary Figs. 20c-d**).

We next assessed reproducibility using Pearson correlation coefficients (PCC). Under absolute quantification, reproducibility was highest within a batch (median PCC: 0.96 for technical replicates) but declined sharply across different platforms, protocols, and laboratories, confirming a strong batch effect (**Fig. 3e**). After ratio-based normalization, median PCC values stabilized (≥ 0.48) across all intra-sample groups, and within each scenario, intra-sample correlation consistently exceeded cross-sample correlation, confirming that batch-specific noise was effectively suppressed (**Fig. 3f**). These results establish ratio-based quantification as a robust foundation for reproducible isoform expression.

Finally, we tested whether this framework could be extended to epitranscriptomics. We profiled NL-methyladenosine (mLA) modifications from raw signal data in two ONT direct RNA batches (**Supplementary Fig. 5**), detecting 25,000 and 60,000 confident mLA sites per sample (**Supplementary Fig. 21**). While absolute mLA levels showed strong within-sample correlation (median PCC ≈ 0.92, **Fig. 3g**), ratio-based quantification again successfully harmonized the data, resulting in clustering by sample types rather than sequencing batch (**Fig. 3h**). This demonstrates that the ratio-based framework can integrate diverse molecular phenotypes, from isoforms expression to RNA modifications.

### Construction and validation of the ratio-based isoform and AS reference datasets

We constructed high-confidence reference datasets for isoforms and AS events using a four-step consensus framework applied to seven high-quality lrRNA-seq batches (SNR ≥ 10) (**Supplementary Fig. 22**)^13,19^. This selection included two ONT direct-RNA, three ONT PCR-cDNA, and two PAB MAS-ISO-Seq batches, leveraging the complementary strengths of both sequencing technologies. We generated separate reference datasets from long-read (LO) and short-read (SO) data. Due to the inherently lower quantitative ambiguity of lrRNA-seq, we selected the LO reference dataset as the primary reference and validated it against the SO dataset using RT-qPCR^24^ (**Supplementary Fig. 23**).

Isoform reference dataset. The LO isoform reference dataset contained 4,588, 2,427, and 3,203 expressed isoforms (ref isoforms) for the D5/D6, F7/D6, and M8/D6 comparisons, respectively, from which we identified 450, 226, and 401 differentially expressed isoforms (ref DEIs) (**Fig. 4a, Supplementary Table 5**). While most reference isoforms were protein-coding, the DEIs were enriched for lncRNA and retained-intron transcripts (**Fig. 4b, Supplementary Table 6**). Orthogonal validation by RT-qPCR showed high concordance with LO-derived logL fold change (logLFC) values (Pearson’s *r* = 0.94, 0.83, and 0.90; RMSE = 0.39, 0.63, and 0.56; **Fig. 4c, Supplementary Table 7**). The dataset also showed high recovery rates for clinically curated MANE transcripts (65.5%–68.2%; **Fig. 4d**) and exceptionally high concordance (Pearson’s *r* > 0.95) with the SO datasets for 7,437 shared isoforms (**Supplementary Figs. 24a-b**), confirming its robustness and reliability.

**Fig. 4.**
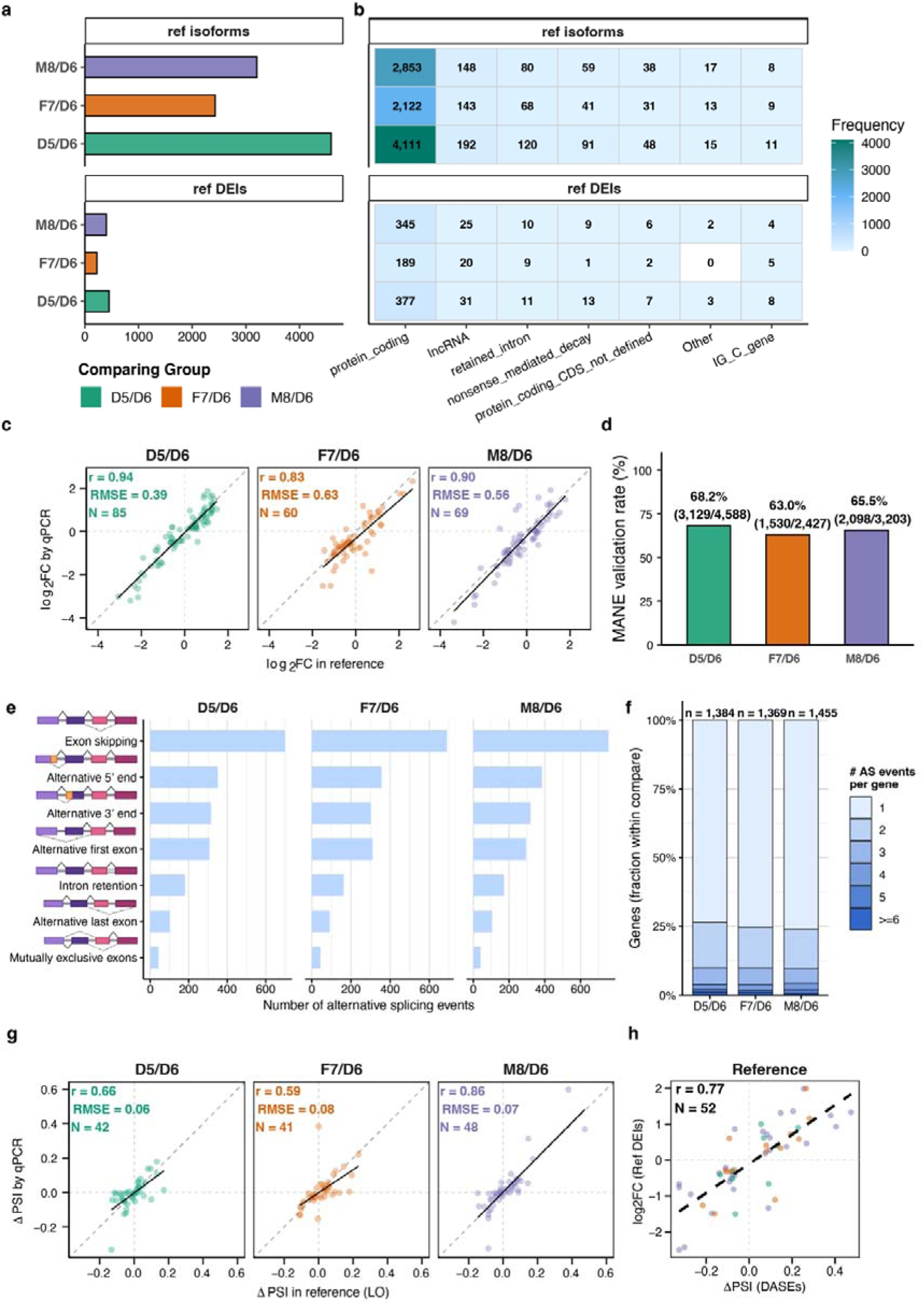
Construction and validation of ratio-based isoform and AS reference datasets. **a,** Numbers of ratio-based isoform reference dataset (ref isoform, upper) and differential isoform expression reference dataset (ref DEI, lower) that pass the consensus filters for each biological contrast (colors: D5 vs. D6 green, F7 vs. D6 orange, M8 vs. D6 purple). **b,** Heatmap tiles report the number of features for each sample pair and biotype category; color intensity encodes frequency. **c,** Orthogonal RT–qPCR validation of reference logL-fold changes for isoforms (one point per isoform; solid line = regression fit, dashed = identity; in-plot text indicates Pearson *r*, RMSE, and N). **d,** Recovery rate of clinically curated MANE transcripts within the LO reference for the three comparisons (values above bars = total MANE isoforms / total reference isoforms). **e,** Summary of alternative splicing (AS) event types identified in the reference datasets across three biological comparisons (D5/D6, F7/D6, and M8/D6). Exon skipping (ES) events were the most abundant. **f,** Distribution of the number of AS events per gene. Most genes exhibited one or two AS events, whereas a small subset showed extensive splicing complexity (> 6 events). **g,** Orthogonal RT–qPCR validation of reference ΔPSI for AS events (one point per AS event; solid line = regression fit, dashed = identity; in-plot text indicates Pearson *r*, RMSE, and N). **h,** Agreement between isoform-level and AS-level differential analyses for 52 matched pairs of differentially expressed features.

AS reference dataset. In the LO AS reference dataset, we identified 2,004, 1,956, and 2,072 AS events for the D5/D6, F7/D6, and M8/D6 comparisons, respectively. Exon skipping (ES) was the most prevalent type, followed by alternative 5′ splice sites (A5) and alternative 3′ splice sites (A3) (**Fig. 4e, Supplementary Table 8**). Splicing complexity was substantial, with >30% of genes containing ≥2 AS events and up to 10% containing ≥4 AS events (**Fig. 4f**). Validation of 41-48 LO-derived AS events (ΔPSI, refDAS) by qPCR confirmed strong agreement with reference ΔPSI values (Pearson’s *r* = 0.66, 0.59, and 0.86; RMSE = 0.06, 0.08, and 0.07, respectively; **Fig. 4g**, **Supplementary Tables 9-10**). Furthermore, we observed a strong global correlation (Pearson’s *r* = 0.77) between logLFC and ΔPSI values for 52 matched DASE-isoform pairs (**Fig. 4h**), demonstrating consistency between isoform-level expression and splicing-level differential analyses. Finally, comparison with the SO AS dataset revealed high concordance (Pearson’s *r* = 0.74) for 4,070 shared AS events (**Supplementary Figs. 25a-b**). These results collectively establish the accuracy and biological validity of the LO isoform and AS reference datasets.

### Hybrid long- and short-read approach improves quantification accuracy

To comprehensively evaluate isoform and AS quantification accuracy, we benchmarked ten analytical pipelines across 34 datasets (12 lrRNA-seq and 22 srRNA-seq) using our validated reference datasets. We compared three fundamental analysis strategies: long-read only (LO), short-read only (SO), and hybrid long- and short-read (LS), assessing performance using relative correlation (RC), root mean square error (RMSE) for quantification, and Matthews correlation coefficient (MCC) for differential feature identification^13^.

The hybrid LS strategy demonstrated uniform superiority across all metrics and datasets. For isoform quantification, LS maintained a high relative correlation (mean LS = 0.92) with the reference datasets, significantly outperforming both LO (0.75) and SO (0.78) strategies (**Fig. 5a**). Crucially, LS reduced quantification error by more than half (mean RMSE = 0.30 vs. 0.76 for LO and 0.60 for SO) and improved differential detection accuracy substantially (mean MCC = 0.78 vs. 0.57 for LO and 0.67 for SO). The performance gains were even pronounced for AS quantification (**Fig. 5b**), where LS boosted the mean relative correlation from 0.39 (LO) and 0.34 (SO) to 0.60, halved the ΔPSI RMSE to ∼0.06 (0.12 for LO and 0.10 for SO), and delivered the highest MCC (0.46 vs. 0.35 for LO and 0.29 for SO). These results indicate that supplementing long-read libraries with matched short-read data provides the optimal balance of accuracy and sensitivity for both isoform and AS event quantification.

**Fig. 5.**
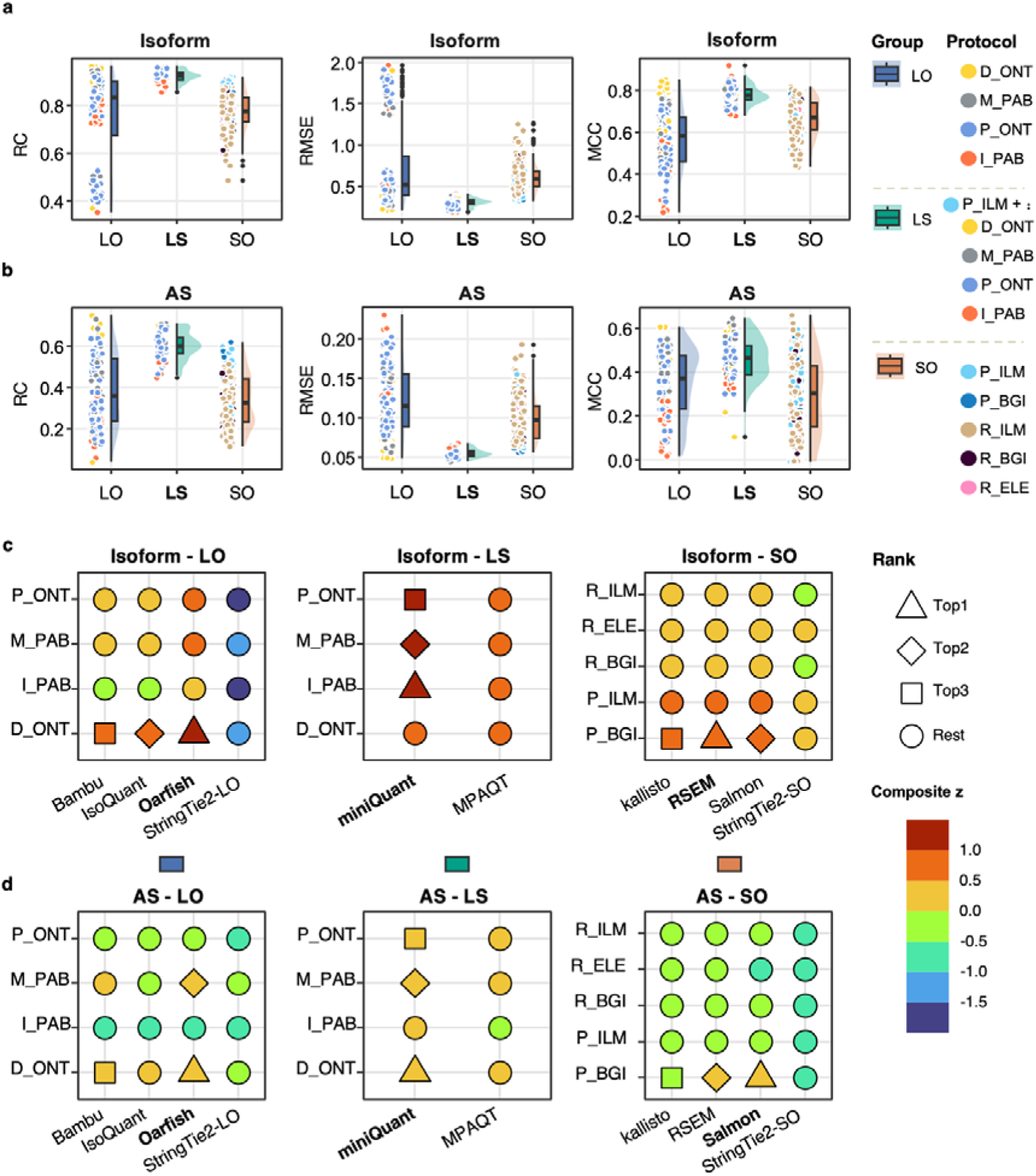
Performance assessment of different sequencing datasets across different strategies. Box-and-jitter plots summarizing Pearson correlation (RC), root mean square error (RMSE; lower is better), and Matthews correlation coefficient (MCC) of every protocol (point) and strategy (box fill: LO = slate-blue, LS = teal, SO = vermillion) for **a,** isoform and **b,** alternative splicing datasets. Ranked bubble plots of the composite z-score that averages RC, sign-reversed RMSE, and MCC of LO, LS, and SO strategies for **c,** isoform and **d,** alternative splicing datasets. The x-axis lists candidate pipelines; the y-axis lists protocols and platforms. Bubble colors encode the composite z-score. Top performers within each facet and strategy are annotated in situ (triangle = Top1, diamond = Top2, square = Top3, circle = Rest.). Pipelines lacking results for a given batch are omitted.

To deconstruct these performance patterns, we ranked all pipeline–batch combinations by a composite z-score incorporating all three metrics. For LO analyses, ONT direct-RNA data processed with Oarfish achieved the highest isoform accuracy, followed closely by IsoQuant and Bambu (**Fig. 5c**), while PacBio MAS-ISO-Seq data analyzed with Oarfish showed strong AS quantification performance (**Fig. 5d**). For SO analyses, Illumina and BGI polyA data quantified by Salmon, kallisto, or RSEM yielded the top performances (**Figs. 5c-d**). Strikingly, incorporating a single short-read library (LS strategy) elevated all batches into the upper performance quartile. The hybrid pipelines, particularly miniQuant, consistently occupied the top ranks (**Figs. 5c-d**), and even shallow-coverage ISO-Seq runs achieved high accuracy through hybrid analysis (**Fig. 5c, Supplementary Fig. 26**).

Collectively, our findings demonstrate that the hybrid LS approach—harnessing the complementary strengths of long-read resolution and short-read depth—achieves the most accurate transcriptome profiling at both isoform and splicing event levels.

### Hybrid long- and short-read assembly extends transcriptome annotation

To further leverage the complementary strengths of lrRNA-seq and srRNA-seq datasets, we performed a hybrid assembly to construct a Quartet-specific reference transcriptome enriched for novel transcript models supported by multiple samples.

We combined seven high-quality long-read batches with one high-quality short-read batch using StringTie3 in hybrid mode. This generated an initial *GENCODE + Quartets* catalogue containing 31,784 novel isoforms and 32,352 novel genes (**Supplementary Fig. 27a**). Applying a stringent filter—requiring detection in at least two samples—yielded a high-confidence set of 120,223 isoforms, comprising 99,420 annotated and 20,803 novel isoforms.

We classified these 20,803 novel isoforms using SQANTI3^29^ based on their relationship to annotated isoforms: 1) 10,570 novel-in-catalog (NIC) isoforms from novel combinations of known splice sites; 2) 7,266 novel-not-in-catalog (NNC) isoforms containing entirely novel splice sites; 3) 1,478 fusion isoforms derived from two or more adjacent genes; and 4) 741 incomplete splice matches (ISM) of 5′ or 3′ fragments of known transcripts (**Supplementary Fig. 27b**).

We next assessed the protein-coding potential of these novel isoforms. ORF (open reading frame) analysis revealed that NIC, NNC, and ISM isoforms were predominantly protein-coding (73%, 56%, and 86%, respectively), whereas other categories were enriched for non-coding or nonsense-mediated decay (NMD)-sensitive transcripts (**Supplementary Fig. 27b**). The hybrid assembly also significantly expanded the annotation of AS events, particularly retained introns, which are under-represented in GENCODE (**Supplementary Fig. 27c**). Although not used for quantification in this study, this curated structural resource provides a foundation for future investigations of transcript diversity in the Quartet standards.

To benchmark the assembly strategy, we compared our hybrid StringTie3 assembly to a long-read-only (LO) assembly generated with Bambu using Gffcompare^30^, which identifies exact matches between chains of introns. Of the 119,334 isoforms discovered by StringTie3, Bambu recovered 88,731 (74.4%). This comparison showed that shared isoform set was enriched for annotated transcripts, while tool-specific isoforms were predominantly novel (**Supplementary Fig. 28a**). Critically, the hybrid StringTie3 approach recovered both a larger total number of isoforms and a higher proportion of NNC isoform—those with entirely novel splice sites—compared to Bambu (**Supplementary Fig. 28b**), underscoring the clear advantage of the hybrid strategy for comprehensive isoform discovery.

## Discussion

In this study, we established the first transcriptome-wide, ratio-based reference datasets for both isoforms and alternative splicing (AS) events using lrRNA-seq, directly addressing the critical lack of a biological ground truth that has hindered the field. Moving beyond earlier benchmarking efforts focused on absolute quantification^10,15,16^, our work provides a comprehensive foundation for future studies through a ratio-based framework. We further demonstrate the unparalleled utility of Quartet reference materials for monitoring and mitigating batch effects across gene expression, isoform quantification, and even epitranscriptomic analyses, underscoring their versatility as a platform for transcriptome standardization.

Our head-to-head benchmarking of ONT and PAB platforms revealed distinct, complementary strengths. The PAB platform offered superior per-base accuracy and read length but introduced quantification bias for short isoforms. Conversely, the ONT platform provided more reliable quantification of sub-kilobase transcripts and, via direct RNA sequencing, uniquely enabled the simultaneous capture of sequence, expression, and m^6^A modification data in a single assay. Furthermore, we consistently found that analytical accuracy declines as resolution increases from gene to isoform to AS-level, highlighting the inherent challenge of high-resolution transcriptomics.

Comparing long- and short-read technologies, we confirmed that lrRNA-seq significantly reduces mapping ambiguity and improves transcript assignment accuracy. Notably, even low-coverage lrRNA-seq reliably detected clinically curated MANE transcripts at a level comparable to deep-coverage srRNA-seq, justifying its use as the backbone of our reference datasets. Critically, ratio-based quantification successfully removed systematic technical offsets, enabling the direct integration and joint analysis of long- and short-read data within a unified quantitative framework for the first time^16^.

The core of our methodological advance is the demonstration that ratio-based quantification effectively eliminates batch effects in multi-center lrRNA-seq studies. While absolute quantification is dominated by technical variance from platforms and pipelines, expressing abundance as sample-to-reference ratios (SRR) sharply attenuated these confounders. This approach restored the visibility of biological signals, significantly enhanced reproducibility across laboratories and protocols, and aligned with emerging best practices in other transcriptomic fields^13,14^, including single-cell^31^ and spatial^32^ applications. We also extended this framework to epitranscriptomics, showing that ratio-based analysis of m6A modification levels from ONT direct RNA sequencing data similarly suppressed batch variability, paving the way for large-scale, integrative studies of RNA modification dynamics^12^.

Leveraging this ratio-based framework, we constructed and rigorously validated the first biological reference datasets for isoform and AS event quantification, with orthogonal RT-qPCR confirmation. These datasets provide an essential resource for benchmarking protocols, developing computational tools, and ultimately supporting the regulatory approval of clinical RNA-seq assays.

A key finding from our benchmarking is the superior performance of the hybrid long- and short-read (LS) strategy^22,23,33^. This approach consistently ranked highest across all performance metrics (RC, RMSE, and MCC)^13,14^, leveraging the structural resolution of long reads with the quantitative precision of short reads. It was also instrumental in building a Quartet-specific transcriptome, enriching annotation with novel isoforms. Thus, integrated LS methodologies represent the optimal path forward for accurate transcriptome analysis. The ratio-based framework provides the foundation for integrating diverse data types, and we encourage the community to build upon this toward a more comprehensive, multi-platform integrative analytic framework.

Nevertheless, our study has limitations. First, the sequencing depth for most lrRNA-seq libraries (∼5 million reads) was modest compared to srRNA-seq, potentially limiting power for low-abundance isoforms. Second, the reference datasets are derived from lymphoblastoid cell lines and thus lack isoforms specific to other tissues or disease states^10^. Third, we did not benchmark the impact of read-alignment algorithms or reference genome/transcriptome versions (e.g. telomere-to-telomere (T2T) genome^34^), which can influence junction detection and isoform identification^29^.

To promote widespread adoption and further methodological development, all raw data, assemblies, ratio-based reference datasets, and Quartet RNA reference materials are publicly available. As long-read technology continues to advance in throughput and accuracy, we anticipate that ratio-based quantification coupled with a hybrid LS strategy will become the new paradigm for reproducible transcriptomics. This framework can be extended to integrate analyses of RNA modifications, fusion transcripts, and RNA edits, uncovering new regulatory mechanisms and therapeutic targets. Furthermore, building a pantranscriptome^35^ based on the T2T genome^36^ using a hybrid LS strategy promises a more complete and accurate representation of the transcriptome.

In conclusion, we have established the first ratio-based reference datasets for full-length transcriptomics, providing a ground truth for benchmarking isoform and AS analysis, RNA modification detection, and other transcriptomic applications. The Quartet RNA reference materials and associated resources constitute an invaluable foundation for methodological development, standardization, and the broader adoption of precise transcriptomics in research and clinical settings.

## Online Methods

### RNA reference materials

All certified Quartet RNA samples (RIN > 9.0) used in this study were obtained from four Epstein–Barr virus–immortalized B-lymphoblastoid cell lines (B-LCLs) derived from the Quartet family and distributed by the Quartet Project^37^. The Quartet Project provides matched multi-omics reference materials, including DNA^38^, RNA^13^, protein^39^, and metabolite^40^ from the same cell-culture batch^12^ under the supervision of Fudan University and the National Institute of Metrology (NIM) of China. RNA reference materials were obtained from GBW09904-GBW09907 (August 22, 2022). The D5/D6/F7/M8 cell lines were provided by a Chinese monozygotic twin quartet, including the father/mother. Quartet RNA reference materials can be requested from the Quartet Data Portal (https://chinese-quartet.org/) or through the National Sharing Platform for Reference Materials (https://www.ncrm.org.cn/Web/Home/EnglishIndex) under the Administrative Regulations of the People’s Republic of China on Human Genetic Resources.

### Library construction and sequencing

According to the Quartet long-read RNA-seq Project study design, 12 tubes of RNA samples were sent to each laboratory, including four groups of the Quartet RNA reference materials with triplicates per group. Library preparation, library quality control, and sequencing were conducted in a fixed order (D5-1, D6-1, F7-1, M8-1, D5-2, D6-2, F7-2, M8-2, D5-3, D6-3, F7-3, and M8-3) in each laboratory to eliminate confounding factors, such as the order of experimental sample processing.

Each laboratory prepared libraries and processed high-throughput RNA-seq. Libraries were constructed using platform-specific methods according to the manufacturer’s protocols for each platform. The libraries were sequenced on long-read instruments, including the PacBio Revio or Sequel IIe and the Nanopore PromethION platforms, in four laboratories. A total of 144 Quartet RNA-seq libraries from 12 batches were finally generated. A single batch of Quartet RNA materials was additionally spiked with SIRV-Set 4 RNA materials^41^, and the libraries were constructed by MAS-ISO-Seq before PacBio sequencing on the Revio platform (**Supplementary Fig. 2**).

### SIRV-SET4 Spike-in ground truth and pipeline validation

SIRV-Set 4^41^ contains 92 ERCC and 69 SIRV transcripts with defined input amounts (**Supplementary Fig. 1**). The ground truths for ERCC and SIRV spike-in transcripts were established based on their concentrations within the samples. The concentrations were taken from the ‘conc (amoles/µl)’ column in the manufacturer’s design table (https://www.lexogen.com/wp-content/uploads/2021/06/SIRV_Set4_Norm_sequence-design-overview_20210507a.xlsx) and rescaled to yield a sum of 10^6^. These values were then treated as the ground truths.

To assess the performance of isoform detection tools in lrRNA-seq, we applied five widely used computational pipelines—IsoQuant (v3.6.1)^4^, StringTie2 (v2.2.1)^42^, Bambu (v3.6.0)^43^, Oarfish (v0.6.2)^28^, and Flair2 (v2.0.0)^44^—to the standardized SIRV-Set 4 datasets. Our comparative analysis indicated that Oarfish, IsoQuant, Bambu, and StringTie2 each offer complementary advantages in sensitivity, precision, and handling of isoforms. Flair2 delivered reliable quantification for the shorter ERCC controls but showed somewhat reduced concordance on the longer SIRV isoforms (**Supplementary Fig. 4**). In addition, we repeatedly encountered technical hurdles when running Flair2’s collapse module across our complete set of 144 libraries, which compromised its consistency in large-scale processing. Consequently, we carried forward Oarfish, IsoQuant, Bambu, and StringTie2 for isoform discovery and quantification.

### Long read RNA-seq processing and quantification

ONT PCR-cDNA pod5 files were basecalled using Dorado with FAST models and converted to FASTQ, and raw FASTQ reads were performed using Pychopper (v2.5.0) and trim_isoseq_polyA (https://github.com/bowhan/trim_isoseq_polyA) to remove adapter sequences and poly-A tails before alignments. ONT direct RNA pod5 files were basecalled using Dorado with SUP models and converted to FASTQ or BAM for alignment. PAB ISO-Seq CCS reads were converted to HiFi reads, while MAS-ISO-Seq reads were preprocessed using the iso-seq toolkit (v4.0.0, https://isoseq.how) according to the standard protocol, at which point reads were ready for alignment.

The clean FASTQ long reads were aligned to the reference genome GRCh38 (ftp://ftp.ncbi.nlm.nih.gov/genomes/all/GCA/000/001/405/GCA_000001405.15_GRCh38/seqs_for_alignment_pipelines.ucsc_ids/GCA_000001405.15_GRCh38_no_alt_analysis_set.fna.gz) using Minimap2 (v2.24-r1122)^45^. The ONT cDNA and direct RNA data were aligned with minimap2 using the parameters “-ax splice -uf -k14”. For the PacBio data, the alignment was performed using “-ax splice:hq -uf” parameters in Minimpa2. Primary alignments with a mapping quality ≥ 40 were extracted with SAMtools^46^ (v1.13, samtools view -q 40 -F 2304), then written to a coordinate-sorted BAM file and indexed for subsequent quantification. Quantification based on the genome-aligned BAM files was performed with IsoQuant (v3.6.1)^4^, Bambu (v3.6.0)^43^ and StringTie2 (v2.2.1)^42^ using the gene model from GENCODE (https://ftp.ebi.ac.uk/pub/databases/gencode/Gencode_human/release_43/gencode.v43. chr_patch_hapl_scaff.annotation.gtf.gz). Additionally, Oarfish (v0.6.2)^28^ was run to quantify transcriptome alignments with model coverage applied, no filters, and otherwise default parameters, followed by the use of tximport (v1.36.1)^6^ to summarize isoform-level and gene-level estimates. All isoform counts were converted to counts per million (CPM), and a pseudocount of 0.01 was added before log2 transformation.

### Protocol comparison in lrRNA-seq

To compare library performance across PacBio and ONT platforms, we applied a unified quality-control workflow. Read-level metrics (total yield, Q-score, and length distribution) were extracted with Nanoplot (v1.42.0)^47^. Mapping statistics were generated after genome alignment: SAMtools (v1.13)^46^ reported overall and primaryLalignment rates, while RSeQC (v4.0.0)^48^ summarized genomic distribution, transcript coverage, and splice-junction detection consistency. All tools were run with default parameters unless otherwise stated.

### RNA modification analysis

Native RNA signal data in POD5 format were first base-called using Dorado (v0.9.6+0949eb8d, rna004_130bps_sup@v5.1.0) to obtain per-base modification probabilities for the mLA DRACH context. The resulting BAM files were summarized using modkit (v0.5.0), which reports, for every covered genomic position, the number of valid reads (Nvalid_cov) and the number classified as modified (Nmod). To ensure high confidence, a site was retained only when (i) Nvalid_cov ≥ 100 and (ii) Nmod ≥ 50^10^. Rows that passed both criteria were collapsed into a unique identifier (chrom_start_end_strand) and taken forward as “detected mLA sites”. The optional percent_modified filter was not applied in this study, but it is recorded for future stringency tuning. For each sample, the retained methylation counts were converted to CPM using edgeR (v4.0.0)^49,50^, and a pseudocount of 0.01 was added before log2 transformation.

### Feature selection

To obtain a technically stable set of isoforms for ratio normalization, we filtered all features in two steps^51^. First, a two-way ANOVA^52^ (factors: batch, sample-type, and their interaction) was applied to each feature; only isoforms showing either no significant terms or a significant sample-type effect without batch or interaction terms (*FDR* < 0.05, Benjamini–Hochberg) were retained. Second, for these candidates, we assessed the reproducibility of reference-vs-target trends across batches by an “angular-concordance” test: batch-wise mean expression profiles (z-scores) of the two sample classes were compared in sliding windows, and the average angle between their fitted slopes was required to fall below a predefined threshold (or to be locally significant when ≥10 batches were available). Features passing both the ANOVA and concordance filters were designated “pass features” and used for ratio normalization.

### Ratio-based quantification

Ratio-based quantification was performed on a feature-by-feature basis within each batch. Specifically, ratio-based quantifications were calculated based on log2CPM values. For each isoform, the mean of expression profiles of replicates of the reference sample(s) (for example, D6) was first calculated and then subtracted from the log2CPM values of that isoform in each study. The same within-batch centring procedure was applied to site-level mLA counts, yielding ratio-based modification estimates that are directly comparable across batches.

### Signal-to-Noise Ratio (SNR)

We quantified discriminative power with the SNR metric previously defined in the Quartet RNA study^13^. Briefly, SNR is the ten-fold logLL of the ratio between (i) the root-mean-square Euclidean distance among biological groups (signal) and (ii) the root-mean-square distance among technical replicates of the same group (noise) in the two-dimensional principal-component (PC1–PC2) space. The calculation was applied to each batch (m = 4 groups, n = 3 replicates per group), using the first two PCs of logL-transformed CPM values. A threshold of SNR ≥ 10 was retained as the criterion for high signal discrimination, and any batch exceeding this cut-off was deemed a high-quality sequencing dataset.

In addition to the SNR values considering all 12 libraries, we iteratively recalculated a leave-one-out “SNR11” (i.e., using any 11 of the 12 libraries) to pinpoint potential within-batch outlier samples.

### Reproducibility of isoform detection and quantification

We used the Jaccard index to quantify the detection reproducibility of isoform or alternative-splicing (AS) calls between two sequencing datasets. It quantifies the overlap between the two datasets by comparing the number of shared isoforms across both datasets. For isoform detection, shared isoforms are defined as those with identical annotated information. The Jaccard Index ranges from 0 to 1, with values closer to 1 indicating a higher level of consistency. The formula is:

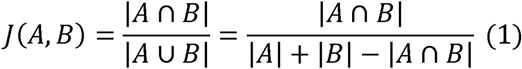

Quantification concordance is used to evaluate the stability of expression levels among technical replicates of the same reference material. It is defined as the Pearson correlation coefficient (PCC) between logL-transformed CPM values from different replicates. By calculating Pearson correlations for each gene, isoform and AS events across technical replicates of the same reference material, we quantify the reproducibility and consistency of the sequencing data.

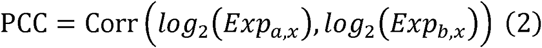

### Short read RNA seq data analysis

The srRNA-seq datasets^13^ were trimmed using fastp (v0.24.1)^53^, removing adapter sequences and low-quality reads. FastQC (v0.12.0) was used for calculating the sequencing depth. The resulting clean reads were then aligned to the reference genome GRCh38 (ftp://ftp.ncbi.nlm.nih.gov/genomes/all/GCA/000/001/405/GCA_000001405.15_GRCh38/seqs_for_alignment_pipelines.ucsc_ids/GCA_000001405.15_GRCh38_no_alt_analysis_set.fna.gz) and reference transcriptome (https://ftp.ebi.ac.uk/pub/databases/gencode/Gencode_human/release_43/gencode.v43. transcripts.fa.gz) using STAR (v2.7.10b)^54^. The generated genome-alignment BAM files were quantified using StringTie2 (v2.2.1)^42^, while the generated transcriptome-alignment BAM files were quantified using RSEM (v1.3.1)^55^. Salmon (v1.10.3)^56^ and kallisto (v0.50.0)^57^ were used for isoform expression quantification in an alignment-free manner. The 50 bootstrap replicates from Salmon were retained to estimate technical variance (over-dispersion) of isoform counts using edgeR^58^. Strand-specific libraries were supplied with the appropriate strandness flags (e.g. --rf in StringTie2, -- rf-stranded in kallisto, -l ISR/FR in Salmon); otherwise, all tools were executed with their default parameters unless stated. Both transcript-level and gene-level estimates were performed and summarized using tximport (v1.36.1)^6^, converted to TPM/CPM, offset with a pseudocount of 0.01, and logL-transformed for downstream analyses.

### Comparison of long- and short-read RNA-seq

Library selection. We restricted the benchmark to libraries whose isoformLlevel signal-to-noise ratio (SNR) was ≥ 10 (**Supplementary Fig. 16 and Supplementary Fig. 19**), retaining seven long-read (lrRNA-seq; ONT direct-RNA, ONT PCR-cDNA and PacBio MAS-ISO-Seq) and five short-read (srRNA-seq; Illumina poly-A, Illumina Ribo-Zero, BGI poly-A, BGI Ribo-Zero and ELE Ribo-Zero) datasets. Comparison aspects include splice-junction coverage per read, read-to-transcript assignments, expression estimate ambiguity, reproducibility of MANE isoform identification and expression concordance between lrRNA-seq and srRNA-seq.

Splice-junction coverage per read. Junction coordinates were extracted from the filtered BAMs with megadepth (v 1.18.0)^59^, a read was classified as junction-spanning if it overlapped every annotated splice junction.

Read-to-transcript assignment. To determine the number of reads that can be uniquely assigned, we processed lrRNA-seq and srRNA-seq data using Bambu (v3.6.0)^43^ without transcript discovery to obtain the read class assignment. We then compared the distribution of the number of transcripts that can be assigned by each read class.

Transcript assignment ambiguity was quantified by overdispersion using the catchSalmon() function in edgeR (v4.0.0) based on the 50 bootstrap replicates of Salmon for srRNA-seq. The catchOarfish() function using identical logic was used to compute the ambiguity based on the 50 bootstrap replicates of Oarfish for lrRNA-seq.

Reproducibility of clinically curated MANE isoforms. Presence/absence of MANE v1.4 transcripts (https://ftp.ncbi.nlm.nih.gov/refseq/MANE/MANE_human/release_1.4/MANE.GRCh3 8.v1.4.summary.txt.gz)^27^ was recorded per library (count > 0). Concordance within and between technologies was measured with the Jaccard index on binary detection matrices.

Expression concordance. Absolute expression consistency was assessed by Pearson correlation (PCC) of logLCPM values across all detected isoforms. To evaluate relative quantification, we computed ratio-based logL-CPM values by subtracting, within each batch, the mean expression of the reference sample (D6) from that of every library, and recalculated the PCC.

### Hybrid-seq data analysis

Isoform quantification using an integrated hybrid long- and short-read approach. We benchmarked two recently released hybrid pipelines. miniQuant (v 1.1)^23^ takes the paired long-/short-read FASTQ files, assigns “community reads” that share splice graphs, and performs a single expectation-maximization (EM) step to produce joint abundance estimates. MPAQT (v 0.3.1)^22^ first computes independent TPMs with Bambu (long reads) and kallisto (short reads) and then fuses the two profiles with its Bayesian EM model. Each long-read library was paired with the corresponding short-read library from the same Illumina batch and processed (i) directly with miniQuant hybrid mode, and (ii) with MPAQT using developer-recommended defaults. Unless strand specificity or library type was required, all tools were run with default parameters.

Quartet-specific transcriptome assembly using a hybrid long- and short-read approach. Seven batches of high-quality lrRNA-seq libraries (SNR ≥ 10) and one batch of matched Illumina SR libraries were assembled with StringTie3 (v3.0.0)^33^ in hybrid mode (--mix), which concatenates the long- and short-read BAMs for each sample. Mono-exonic (unspliced) contigs were excluded. Each sample’s isoform assembly was then merged using StringTie3 –merge, and each isoform was re-quantified for each sample with StringTie3 –estimate against GENCODE v43. Isoforms were compared with the GENCODE v43 reference annotation to determine novelty. All annotated isoforms were then kept first, while novel isoforms were kept only if they were expressed in ≥ 3 samples within a batch. The combined set was refined using SQANTI3 (v5.2.0)^29^ with default quality-control filters and the same GENCODE annotation. Finally, any isoform failing SQANTI3 flags or falling below the re-quantified expression threshold was discarded, yielding the Quartet hybrid reference transcriptome.

### Differentially expressed features (DEFs)

Differential expression analyses were implemented using the edgeR (v4.0.0)^49^ according to guidelines from the package. An isoform or gene was considered differentially expressed in a batch between two sample groups if two-sided *P* < 0.05 and |log2 FC| ≥ 1 using the edgeR package for upregulation or downregulation, respectively.

### Identification and quantification of alternative splicing events and DASEs

Alternative splicing was assessed with a unified SUPPA2 (v2.4)^60^ workflow. First, the GENCODE reference was decomposed into seven canonical event classes using SUPPA2 prepareEvents, including skipping exon (SE), alternative 5’/3’ splice sites (A5/A3), mutually exclusive exons (MX), retained intron (RI), and alternative first/last exons (AF/AL). Next, sample-specific transcript TPMs were supplied to SUPPA2 quantifyPSI to obtain percent-spliced-in (PSI) values for every event. Differential AS events (DASEs) were called with SUPPA2 diffSplice; events with |ΔPSI| ≥ 0.05 and *P* < 0.05 were retained.

### Reference datasets construction

We constructed the ratio-based isoform and AS reference datasets using a four-step consensus framework (**Supplementary Fig. 22**), modified from our previous study^13,19^. First, within each technology (long- or short-read), we performed per-sample quantification and applied intra-batch (detected in ≥ 50% of replicates) and inter-batch (detected across all batches) filters to retain only robustly observed isoforms and AS events. Second, for each key pairwise comparison (D5/D6, F7/D6 and M8/D6), we computed relative expression and identified putative differential expression isoforms (DEIs) with edgeR (v4.0.0)^49^ (*P* < 0.05 and |log2 FC| > 1 for isoforms), and differential alternative splicing events (DASEs) with SUPPA2 (*P* < 0.05 and |ΔPSI| ≥ 0.05 for AS events). Third, we required that each candidate feature be detected in every sample pair and supported by at least half of all quantification pipelines, summarizing effect sizes as mean log_2_FC or ΔPSI and median *P*-value across datasets. Finally, platform-specific reference sets were generated by consensus voting—retaining events observed by ≥ 50% of pipelines.

Independent reference datasets were generated separately from the long-read and short-read libraries, enabling a direct, technology-specific performance comparison.

### qPCR validation

The qPCR validation protocol was implemented using complementary approaches: dye methods for initial screening and probe methods to resolve discordant results. Primer sets targeting 121 isoforms and 59 AS events for three sample pairs and corresponding TaqMan probes were designed using the BeaconDesigner tool. Like previous studies^13,14^, *C1ORF43* was also used as the reference gene in this study. The primers and probes were synthesized by Sangon Biotech, and their detailed sequence information is shown in **Supplementary Tables 11-14**.

The RT-qPCR reaction was performed in two steps. First, the reverse transcription reaction was performed by mixing 1 µl of RNA with Buffer, Enzyme, Oligo dT Primer, Random 6 Primer, and nuclease-free water to a final volume of 10 µl (Takara, No. RR037A). The reaction was incubated at 37°C for 15 min, followed by incubation at 85°C for 5 s, and finally terminated at 4°C. Next, qPCR was performed using the above cDNA as a template. For either the dye or probe method, the reaction system consisted of 1 µl of cDNA template and 19 µl of reaction system in a total volume of 20 µl (dye reagent is Takara, No. RR820A; probe reagent is Takara, No. RR390A). qPCR was performed on an Applied Biotechnology 7500 Real-Time PCR system with an initial denaturation temperature of 95°C for 30 seconds, followed by 40 cycles (95°C for 5 seconds and 55°C for 34 seconds). Three technical replicates were performed for each sample and gene.

Relative fold changes of samples compared to D5/D6, F7/D6, and M8/D6 were calculated using the ΔΔCt method, with *C1ORF43* as the endogenous control. Differential expression was determined in a two-step manner: a Student’s t-test *P* < 0.05 was first required, followed by applying biological thresholds. For isoforms, a |log2 FC| ≥1 threshold was considered significant; for alternative splicing (AS) events, a ΔPSI ≥ 0.05 or ≤ –0.05 threshold was used.

### Reference-based performance metrics

To benchmark each RNA-seq workflow against the Quartet ratio reference, we computed three orthogonal, reference-anchored statistics^13,14^:

### Relative Correlation (RC)

Relative Correlation (RC) was measured by Pearson’s correlation coefficient (PCC) to evaluate the accuracy of isoform and AS quantification. This involved examining the correlation between sample-pair relative expression in the test datasets, qPCR datasets, and the Quartet reference datasets for assessing relative isoform and AS expression.

### Root Mean Square Error (RMSE)

Root Mean Square Error (RMSE) was used to measure the absolute deviation from Quartet reference datasets, implemented with the rmse() function in the Metrics R package (v 0.1.4).

### Matthews Correlation Coefficient (MCC)

Matthews Correlation Coefficient (MCC), a robust bioinformatics metric that encapsulates both sensitivity and specificity, was employed to assess the concordance between DEIs and DASEs detected from our test datasets and Quartet reference datasets. This approach amalgamates true-positive and true-negative calls through a consensus voting mechanism. By discerning DEFs and non-DEFs, we determined the respective counts of true positives (TP), true negatives (TN), false positives (FP), and false negatives (FN), thereby enabling the calculation of MCC.

The MCC is defined as:

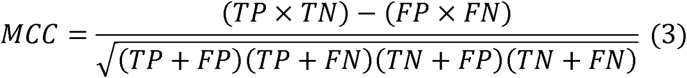

where TP, TN, FP, and FN represent true positives, true negatives, false positives, and false negatives, respectively.

### Composite Z-score

For each library, we report RC, RMSE, and MCC separately at the isoform and AS levels. After reversing the sign of RMSE (so that higher values denote better performance) and z-standardizing each metric, we take the arithmetic mean of the three normalised scores to yield a single composite performance metric used for downstream ranking.

### Statistical analysis

Analyses used R 4.5.0 (https://www.r-project.org). PCA was conducted with unit variance scaling, using the prcomp function. Hierarchical clustering analysis (HCA) was performed using the R package pheatmap version 1.0.13 (https://rdrr.io/cran/pheatmap/). Data visualization was implemented using the R package ggplot2 version 3.5.2 (https://ggplot2.tidyverse.org).

## Data availability

The raw sequence data used in this paper have been deposited in the Genome Sequence Archive (GSA) (under accession code of GSA-Human: PRJCA039455, https://ngdc.cncb.ac.cn/gsa-human/s/cq844JS3) of the China National Center for Bioinformation. The raw sequence data are also available in NODE (under accession code: OEP00006163, https://www.biosino.org/node/review/detail/OEV00000650?code=MAYPZ6K2). Moreover, we have uploaded the expression data and reference datasets to the Quartet Data Portal (http://chinese-quartet.org) for the community to conveniently access and share. The expression matrices of isoforms, and AS events can be accessed at figshare (https://figshare.com/projects/A_ratio-based_framework_using_Quartet_reference_materials_for_integrating_long-_and_short-read_RNA-seq_data/262438).

## Code availability

The source codes for upstream analytical pipelines and the data visualization in this manuscript are available at GitHub: https://github.com/qingwangch/quartet_rna_reference.

## Acknowledgements

We thank the Quartet Project team members who contributed their time and resources to the design and implementation of this ambitious project. This study was supported in part by the National Key R&D Project of China (2023YFC3402501), the National Natural Science Foundation of China (T2425013, 32370701, 32470692, 32400539, and 32170657), the Natural Science Foundation of Shanghai (24JS2840100), Shanghai Municipal Science and Technology Major Project (2023SHZDZX02), the Beijing Natural Science Foundation (L254025), and the 111 Project (B13016). We are grateful to computing resources provided by WEHI HPC (Milton), CFFF (Computing for the Future at Fudan), and the Human Phenome Data Center of Fudan University. This work was also supported by a grant of the China Scholarship Council (202406100193). Fig.1, Supplementary Fig. 2, Supplementary Fig. 5, Supplementary Fig. 22 and Supplementary Fig. 23 in this paper were created with BioRender (https://biorender.com).

## Competing interests

X.L. and H.C. are employees of Wuhan Benagen Technology Co., Ltd. The other authors declare no competing financial interests.

## Authors’ contributions

Y.Z., L.S., R.Z., and M.E.R. conceived and supervised the study. Q.C., Y.L., W.H., X.L., H.C. and L.D. coordinated and/or performed NGS library preparation and sequencing. Q.C., X.G., Y.M., and S.D. performed data analysis and/or interpretation. D.W. and J.Z. performed the qPCR validation experiments. Q.C. managed the datasets and generated most figures. Y.X. performed the RNA modification analysis. Q.C. and X.G. wrote the initial draft. Y.P.Y., D.W., Y.T.Z., Y.Y., J.L., R.Z., Y.Z., M.E.R, and L.S. critically revised the manuscript. All authors reviewed and approved the final manuscript. Quartet Project participants generously contributed time and resources essential to the completion and analysis of this study.

## Notes

https://figshare.com/projects/A_ratio-based_framework_using_Quartet_reference_materials_for_integrating_long-_and_short-read_RNA-seq_data/262438

