## Supplementary figures for "A ratio-based framework using Quartet reference materials for integrating long- and short-read RNA-seq"

### Supplementary Fig. 1. Isoform, abundance, and length complexity in SIRV-Set4 RNA materials.

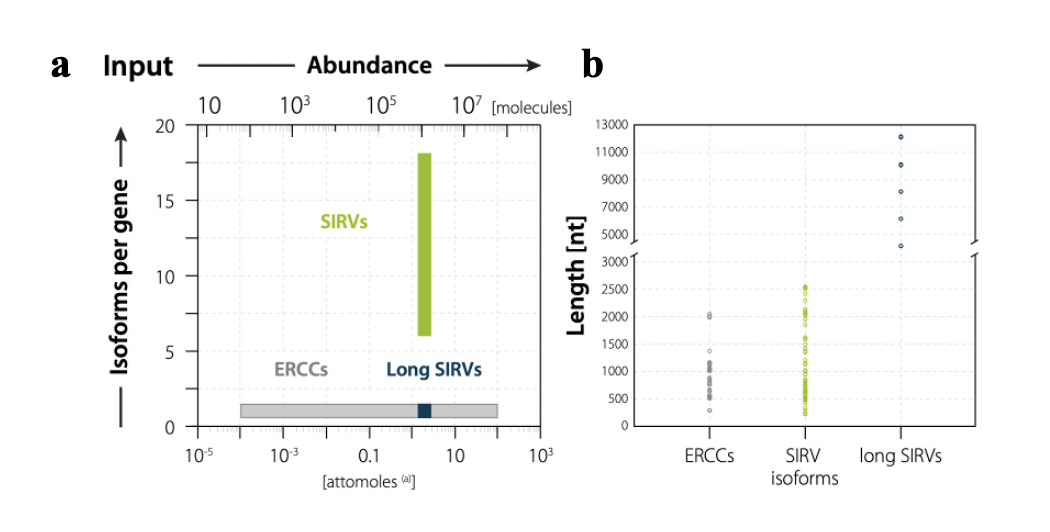

**Supplementary Fig. 1** Isoform, abundance, and length complexity in SIRV-Set4 RNA materials.

**a**, Isoform and abundance complexity and **b**, Length complexity of ERCC and SIRV isoforms in SIRV-Set4 RNA.

### Supplementary Fig. 2. Protocol for spiking SIRV-Set4 into Quartet RNA samples.

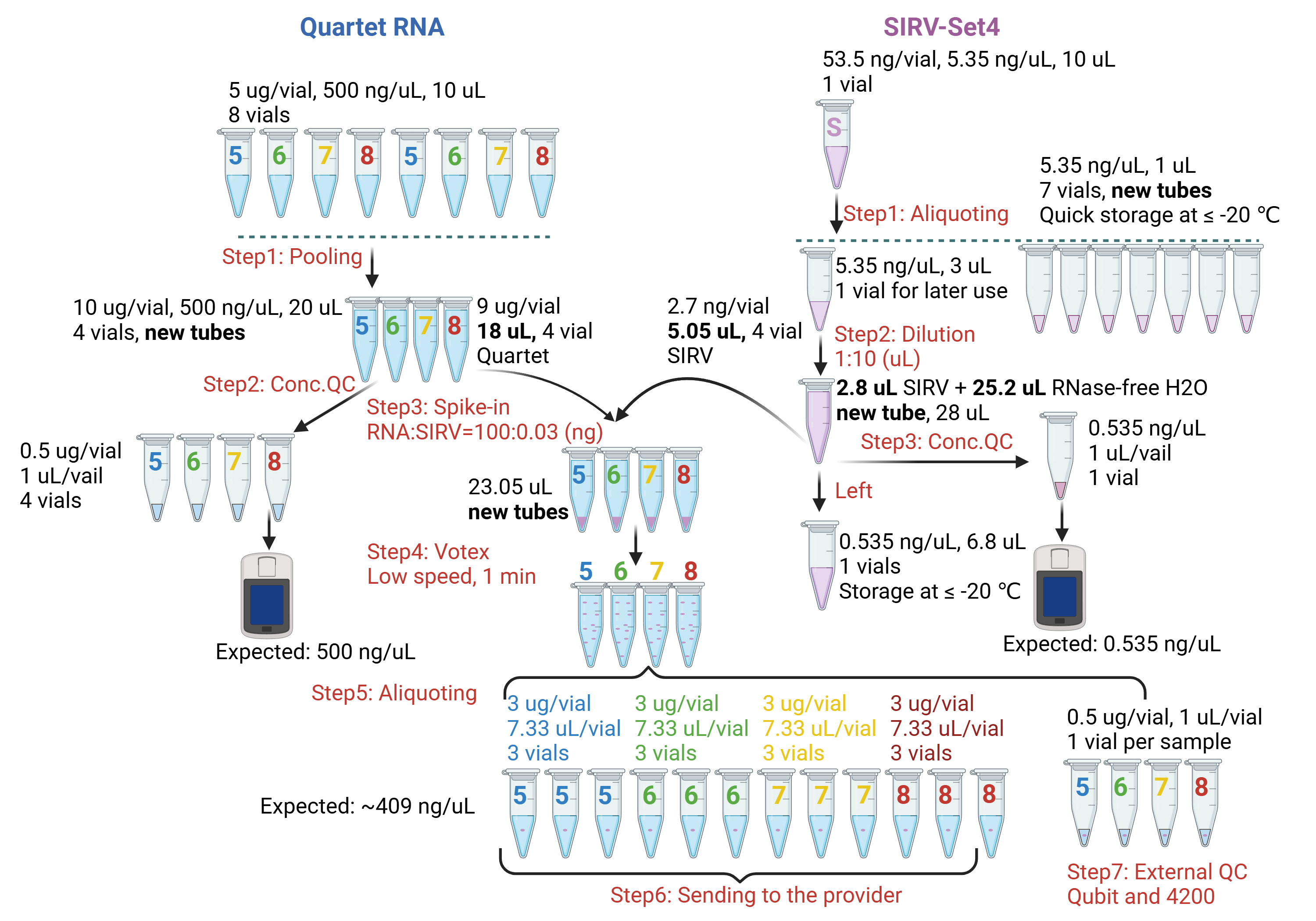

**Supplementary Fig. 2.** Protocol for spiking SIRV-Set4 into Quartet RNA samples.

### Supplementary Fig. 3. Validation of analysis pipelines based on SIRV-Set4 RNA materials.

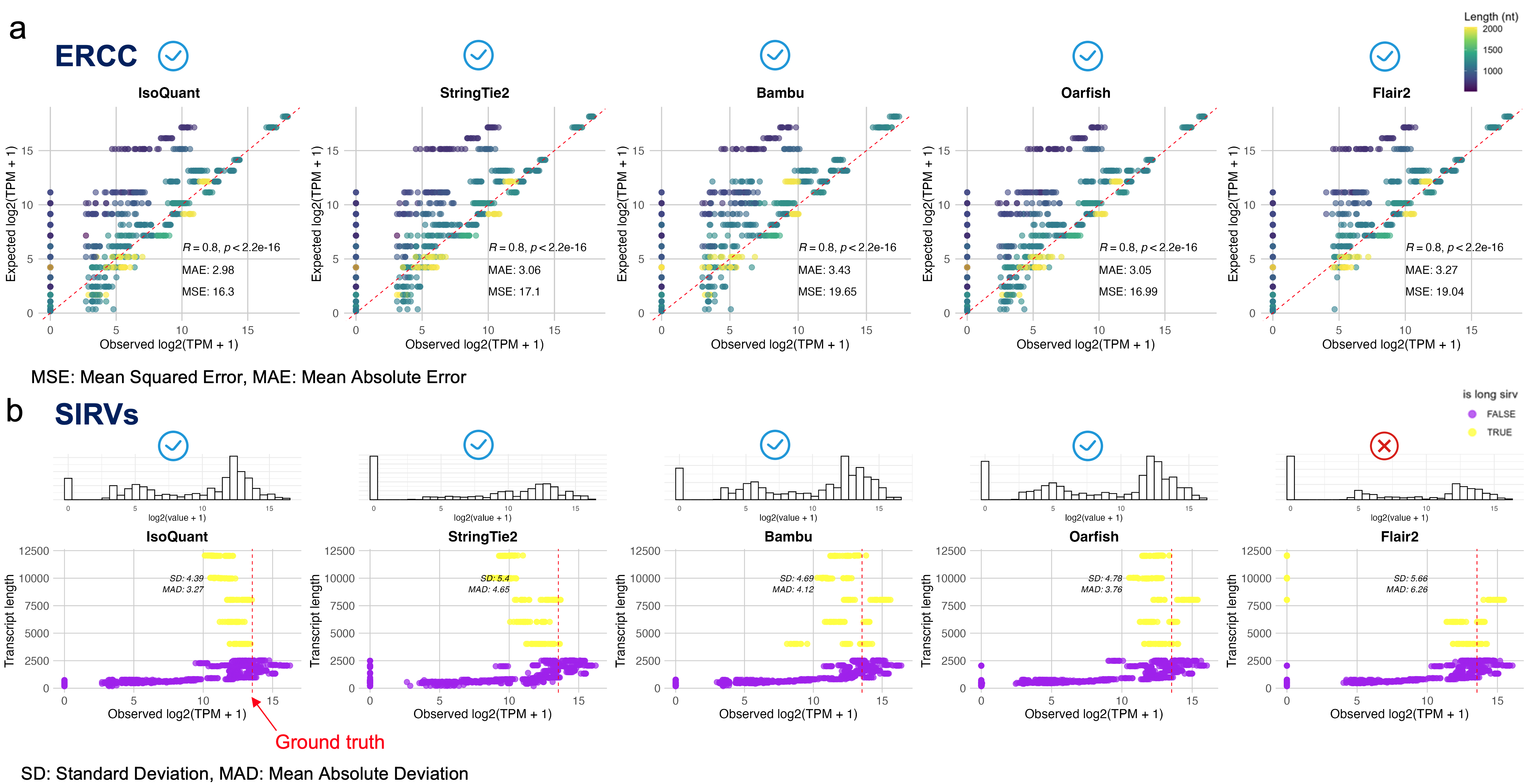

**Supplementary Fig. 3.** Validation of analysis pipelines based on SIRV-Set4 RNA materials.

**a**, ERCC transcripts. For each long-read caller (IsoQuant, StringTie2, Bambu, Oarfish, Flair2), the expected log2(TPM+1) of 92 ERCC controls (y-axis) is plotted against the observed log2(TPM+1) (x-axis). Points are colored by transcript length (legend, top right). The dashed red diagonal indicates perfect concordance. Pearson’s R, mean absolute error (MAE), and mean squared error (MSE) are reported in each panel.

**b**, SIRV transcripts. For the same five callers, scatter plots compare observed versus ground-truth log2(TPM+1) for 69 SIRV isoforms (purple = short SIRV isoforms, yellow = long SIRV isoforms). The marginal histogram above each plot shows the distribution of observed expression. Dashed orange lines denote the ground-truth abundance used during the spike-in titration. Length dependence is visualized by vertical positioning of points (longer isoforms higher). Together, the ERCC and SIRV assessments demonstrate that IsoQuant, StringTie2, Bambu, and Oarfish achieve high quantitative accuracy, whereas Flair2 exhibits systematic underperformance for long SIRVs and is excluded from downstream analyses. Blue checkmarks denote tools that will be carried forward into further analyses, whereas the red cross identifies a tool that will be excluded.

### Supplementary Fig. 4. Overview of publicly available long-read RNA-seq resources.

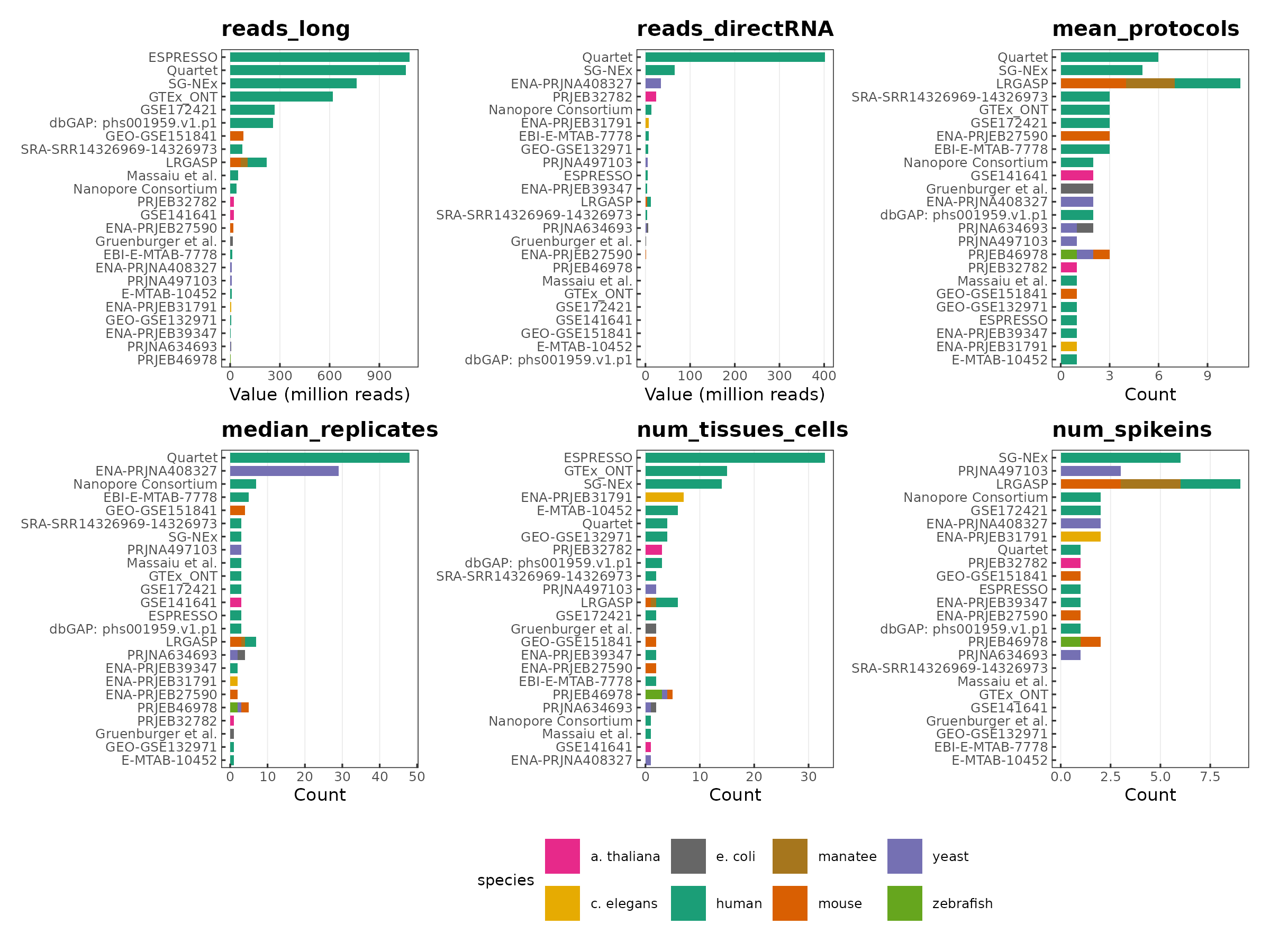

**Supplementary Fig. 4.** Overview of publicly available long-read RNA-seq resources.

Horizontal bar charts rank 24 public or consortium datasets by six key attributes (the largest value at the top of each panel). Colors indicate the organism sequenced (legend, bottom).

### Supplementary Fig. 5. Overview of sequencing data analysis pipelines.

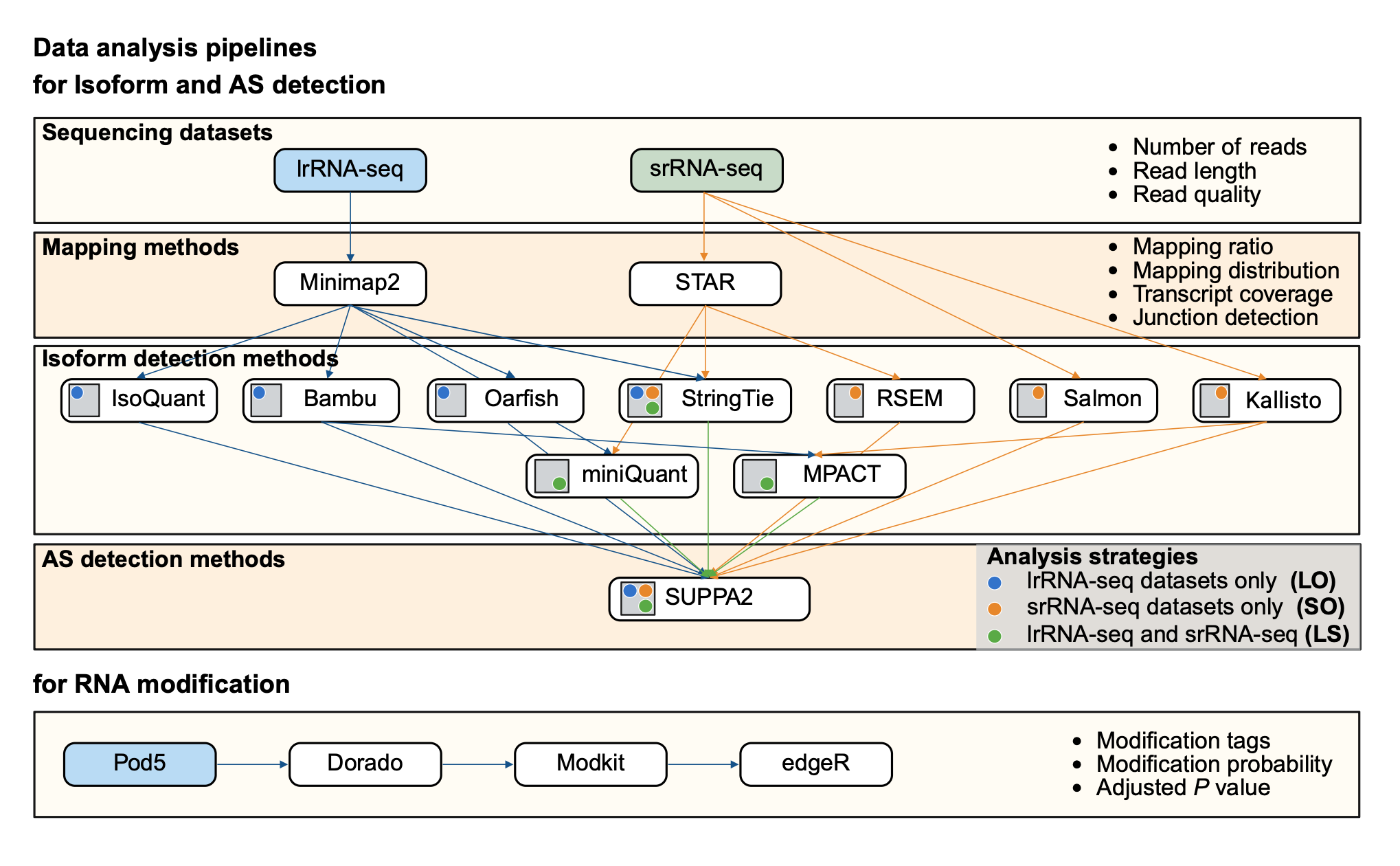

**Supplementary Fig. 5.** Overview of sequencing data analysis pipelines.

Top, isoform and alternative splicing (AS) detection workflows. Long-read RNA-seq (lrRNA-seq) and short-read RNA-seq (srRNA-seq) datasets are mapped with *Minimap2* and *STAR*, respectively; mapping statistics (ratio, distribution, coverage, and junction detection) are recorded. Isoform reconstruction/quantification is performed with long-read callers (*IsoQuant*, *Bambu*, *Oarfish*), short-read callers (*StringTie*, *RSEM*, *Salmon*, *Kallisto*), and hybrid quantifiers (*miniQuant*, *MPACT*). Isoform expressions are passed to SUPPA2 for AS detection. Colored circles denote the three analysis strategies: lrRNA-seq only (LO, blue), srRNA-seq only (SO, orange), and combined long + short (LS, green). Bottom, RNA-modification workflow. Raw pod5 files are base-called with Dorado, annotated with Modkit, and tested with edgeR to yield per-site modification tags, probabilities, and adjusted *P* values. Right-hand labels list the principal evaluation metrics for each tier.

### Supplementary Fig. 6. Raw long-read sequencing statistics across different batches.

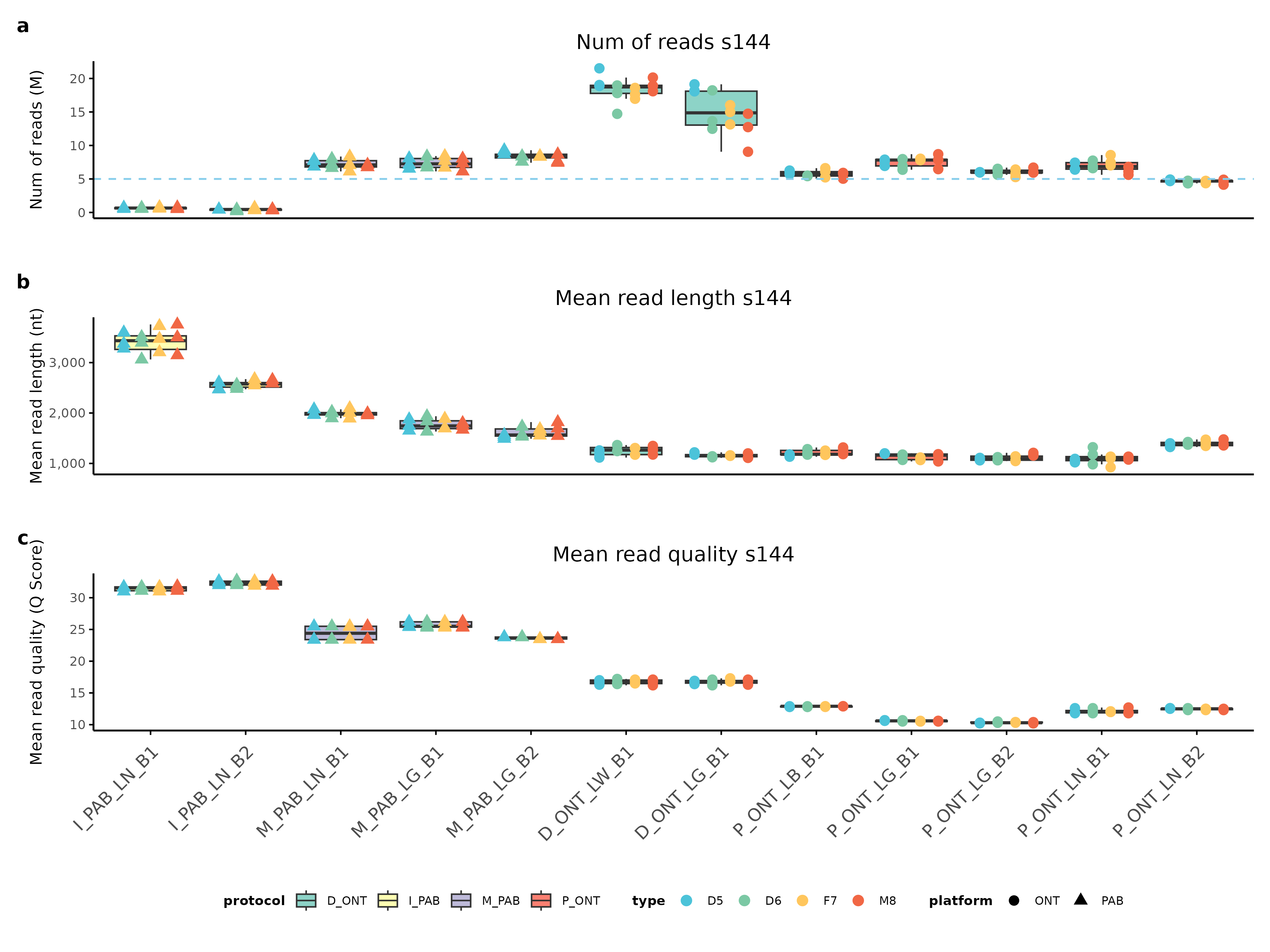

**Supplementary Fig. 6.** Raw long-read sequencing statistics across different batches.

**a**, Total read count per batch (millions); the horizontal dashed line marks the 5 M-read threshold for downstream analysis. **b**, Mean read length and **c**, mean read quality (Phred Q-scores) for the same batches. Boxplots summarize replicate libraries, colored by library-prep protocol (D_ONT, I_PAB, M_PAB, P_ONT), while overlaid points are individual replicates; point color encodes sample type (D5, D6, F7, M8), and point shape distinguishes sequencing platforms (black circle = Oxford Nanopore, black triangle = PacBio; “s144” indicates a collection of 144 samples).

### Supplementary Fig. 7. Mapping ratio across different batches.

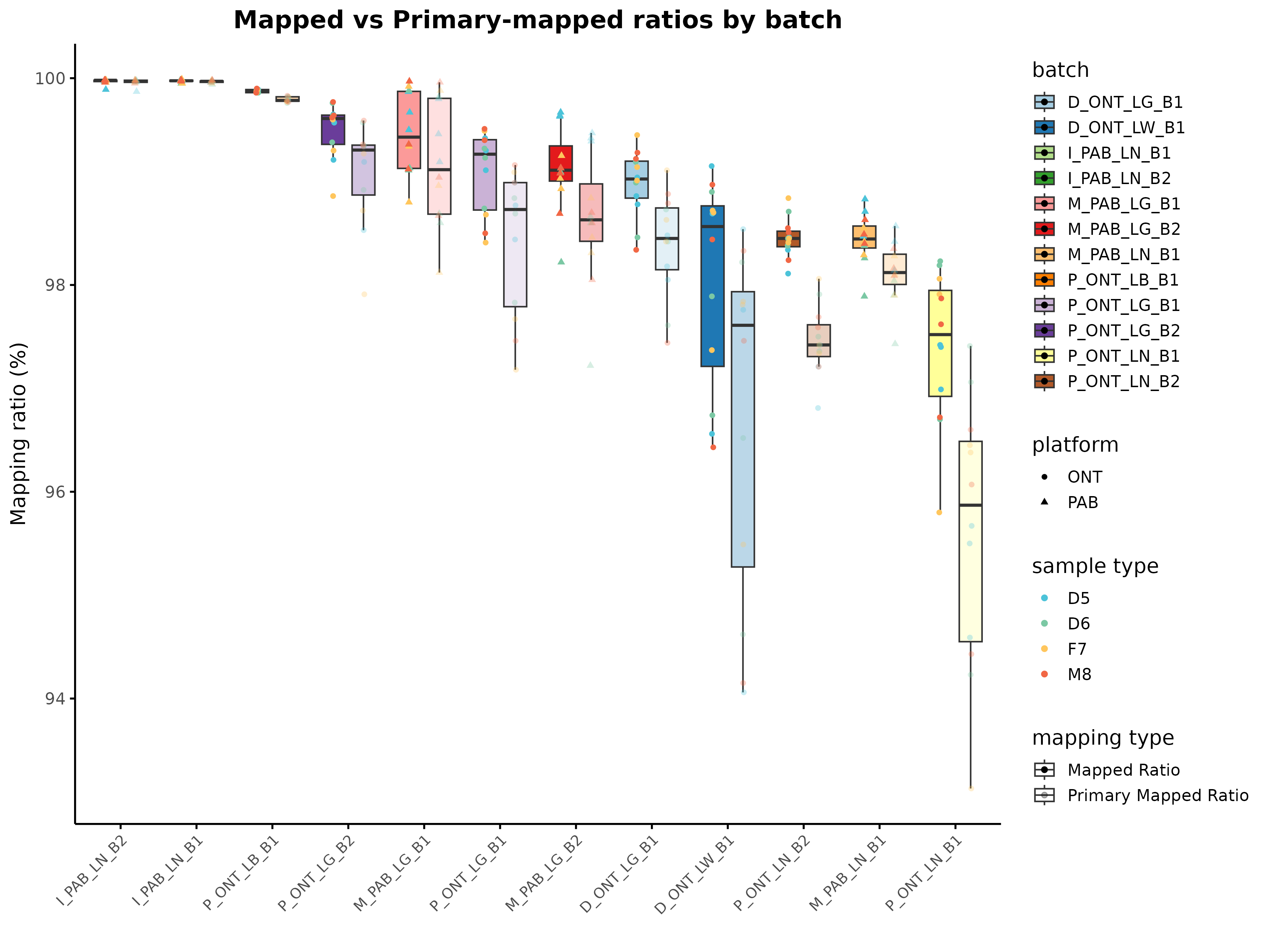

**Supplementary Fig. 7.** Mapping ratio across different batches.

Bar plots show the percentage of reads that align to the reference genome. Boxes are filled by batch color; solid outlines represent the overall mapped ratio, while dashed outlines indicate the primary mapped ratio. Overlaid points are individual replicate libraries: point shape distinguishes the sequencing platform (black circle = Oxford Nanopore, black triangle = PacBio), and point color marks the sample type (D5, D6, F7, or M8).

### Supplementary Fig. 8. Mapping distribution across different batches.

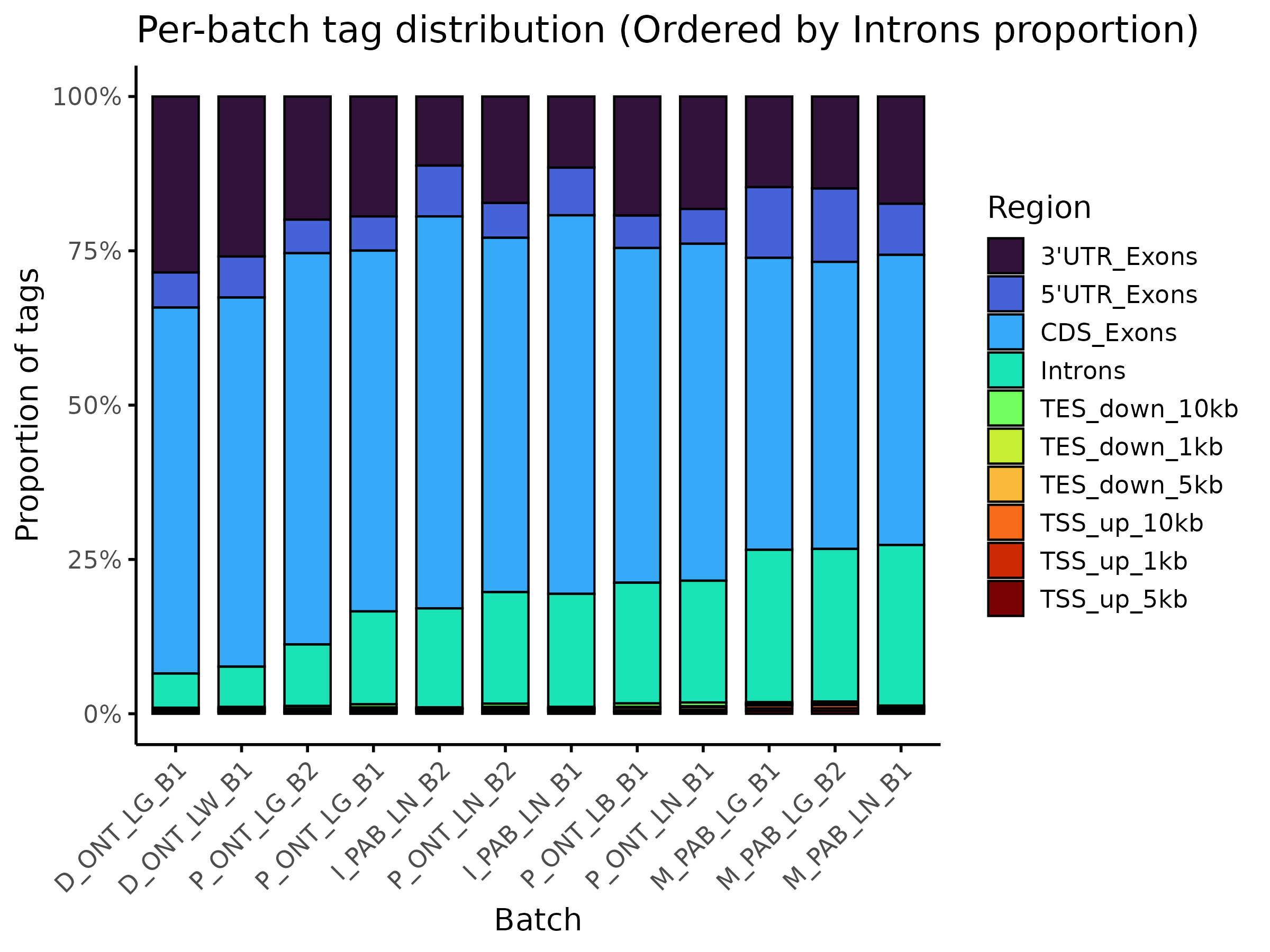

**Supplementary Fig. 8.** Mapping distribution across different batches.

Stacked bars show the fraction of mapped tags that originate from ten genomic regions, with batches ordered by decreasing intronic content. Colors denote annotation categories: 3′-UTR exons (dark purple), 5′-UTR exons (indigo), coding exons (cyan), introns (teal), downstream of the transcription-end site (TES_down 1 kb, 5 kb, 10 kb; light-to-dark greens), and upstream of the transcription-start site (TSS_up 1 kb, 5 kb, 10 kb; light-to-dark reds).

### Supplementary Fig. 9. 5′/3′ coverage bias of long-read libraries across batches.

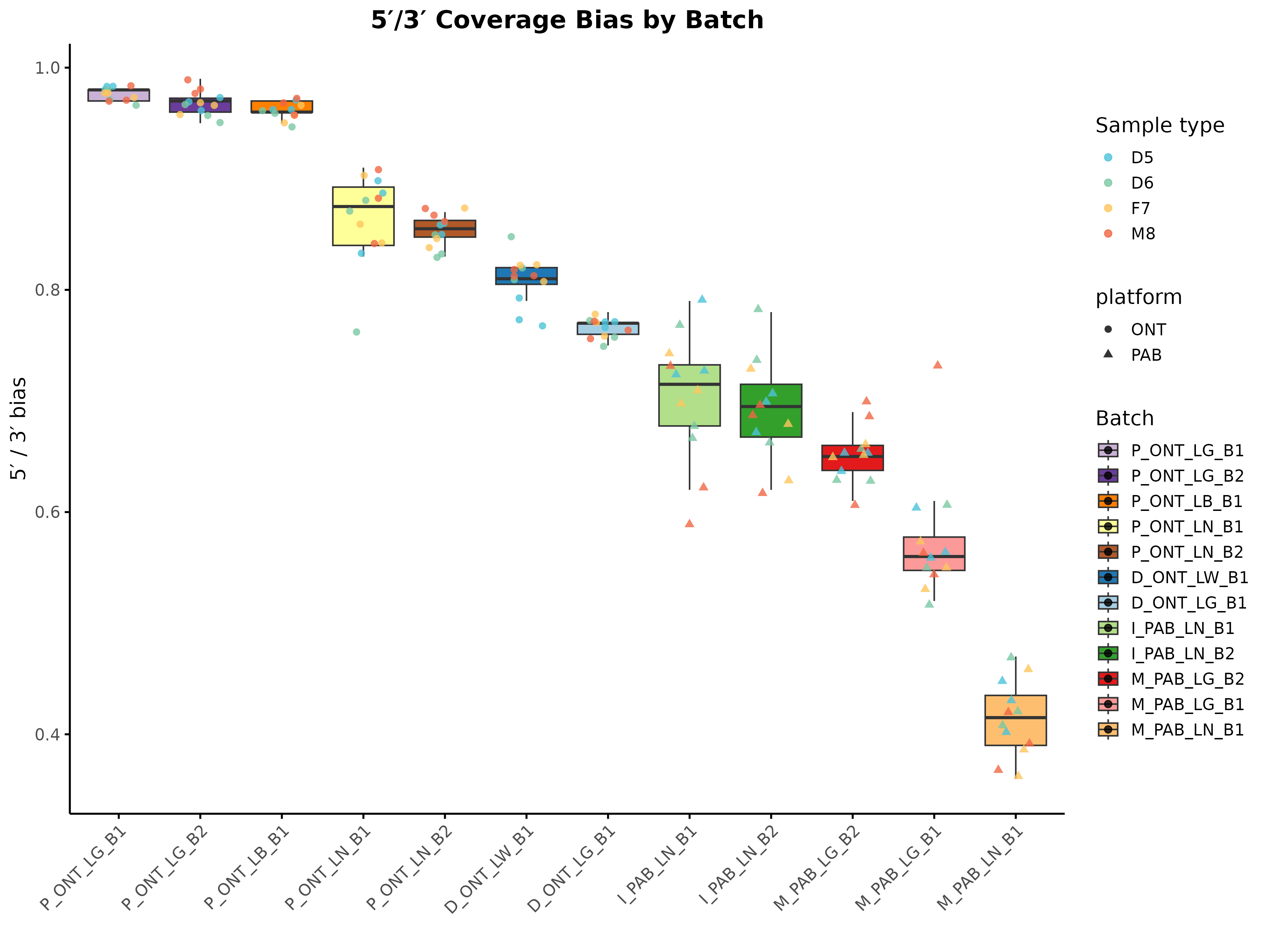

**Supplementary Fig. 9.** 5′/3′ coverage bias of long-read libraries across batches.

The 5′/3′ bias is calculated as the ratio of median read depth at the 5′ end to that at the 3′ end of each transcript (a value of 1 indicates perfectly uniform coverage). Boxplots summarize replicate libraries for 12 batches, ordered left-to-right from lowest to highest bias. Batches are color-coded for ease of comparison. Overlaid points represent individual libraries: point shape denotes the sequencing platform (black circle = Oxford Nanopore, black triangle = PacBio), and point color indicates the sample type (D5, D6, F7, M8).

### Supplementary Fig. 10. Consistency of splice-junction detection across batches.

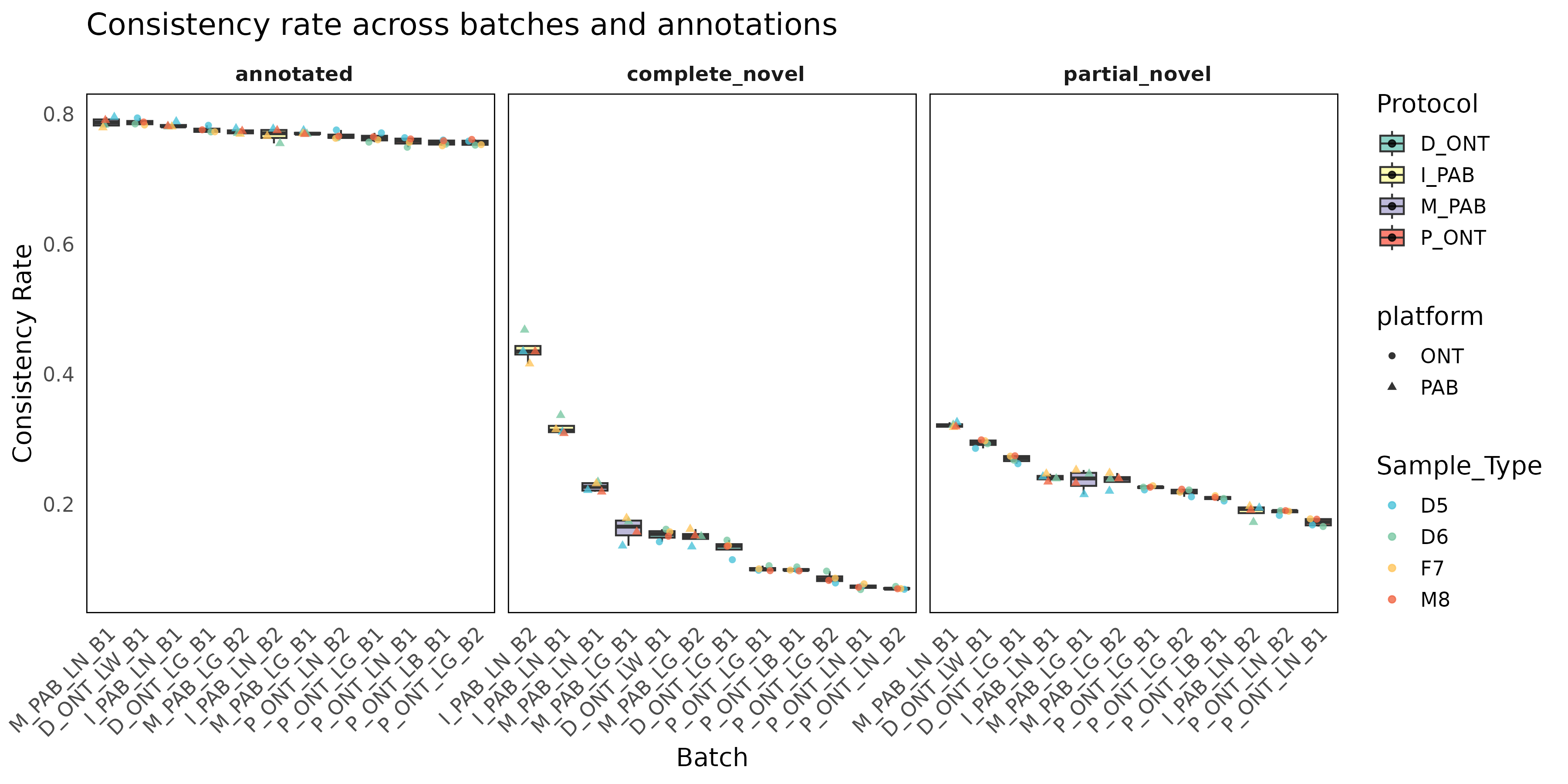

**Supplementary Fig. 10.** Consistency of splice-junction detection across batches.

Box plots depict the per-replicate consistency rate—the fraction of junctions recovered in two independent alignments of the same library—for three annotation classes: previously annotated junctions, complete-novel junctions (both donor and acceptor unannotated), and partial-novel junctions (one annotated splice site). Batches (x-axis) are ordered within each panel by decreasing median consistency. Box fill indicates the library-prep protocol (D_ONT, I_PAB, M_PAB, P_ONT), point shape denotes sequencing platform (black circle = Oxford Nanopore, black triangle = PacBio), and point color marks the sample type (D5, D6, F7, M8).

### Supplementary Fig. 11. Gene, isoform, and AS detection across batches.

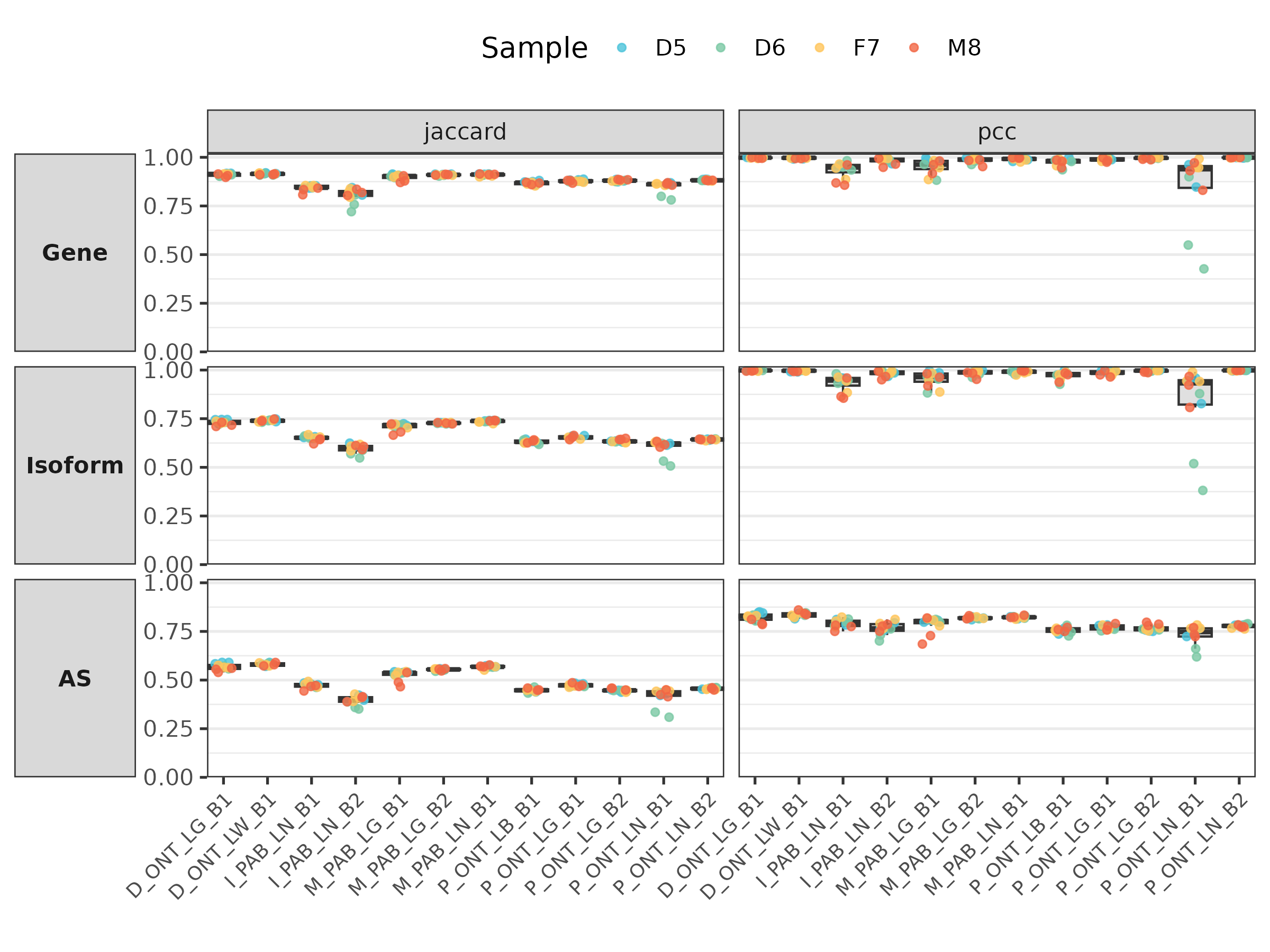

**Supplementary Fig. 11.** Gene, isoform, and AS detection across batches.

For each of the 12 long-read batches, we calculated—across the four biological replicates (colored dots: D5 cyan, D6 green, F7 gold, M8 coral)—(i) the Jaccard index of feature presence/absence and (ii) the Pearson correlation coefficient (PCC) of log₂CPM values. Results are shown side-by-side for the gene, isoform, and alternative-splicing (AS) layers (rows). Boxes mark replicates whose values fall outside the interquartile range of their batch, flagging potential technical outliers. Concordance is highest at the gene level and drops progressively for isoforms and AS events, underscoring the greater reproducibility challenge as analytical resolution deepens.

### Supplementary Fig. 12. Signal-to-noise ratios for each long-read batch at three analytical layers.

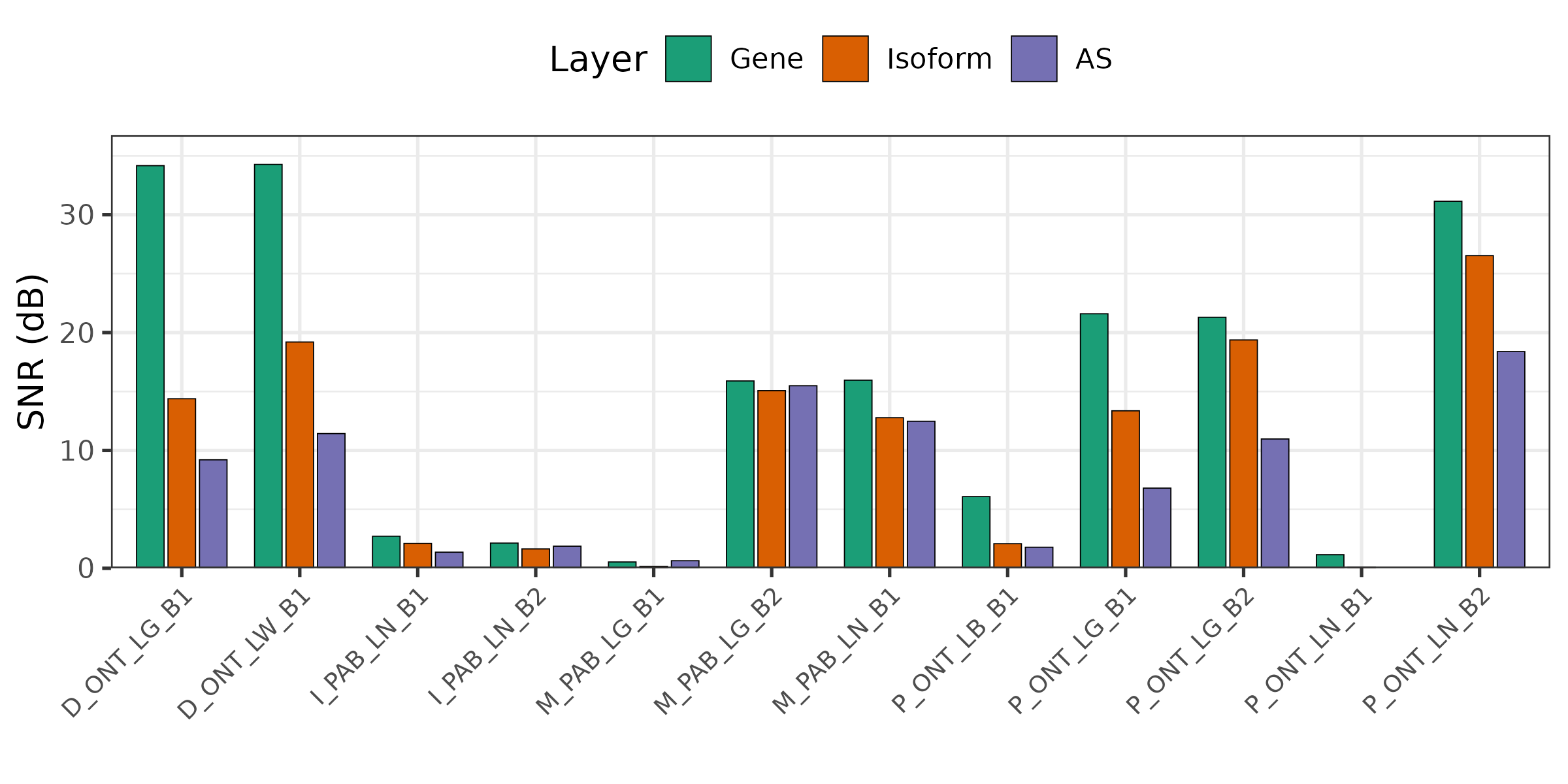

**Supplementary Fig. 12.** Signal-to-noise ratios for each long-read batch at three analytical layers.

For the twelve lrRNA-seq libraries, SNR was computed separately at the gene, isoform, and alternative-splicing (AS) levels (bars colored green, orange, and purple, respectively). Height indicates the decibel SNR derived from pair-wise Euclidean distances in the first two PCA dimensions (Methods). ONT direct-RNA batches (D_ONT_LG_B1, D_ONT_LW_B1) show the strongest discrimination (SNR > 30 dB at the gene layer), whereas MAS-ISO-Seq (M_PAB_LG/LN) and PacBio ISO-Seq (I_PAB_LN) display progressively lower values. Across all protocols, SNR decreases from genes to isoforms and again to AS events, illustrating the rising analytical difficulty with increasing transcriptomic resolution.

### Supplementary Fig. 13. SNR across datasets based on the SNR and SNR11 criterion.

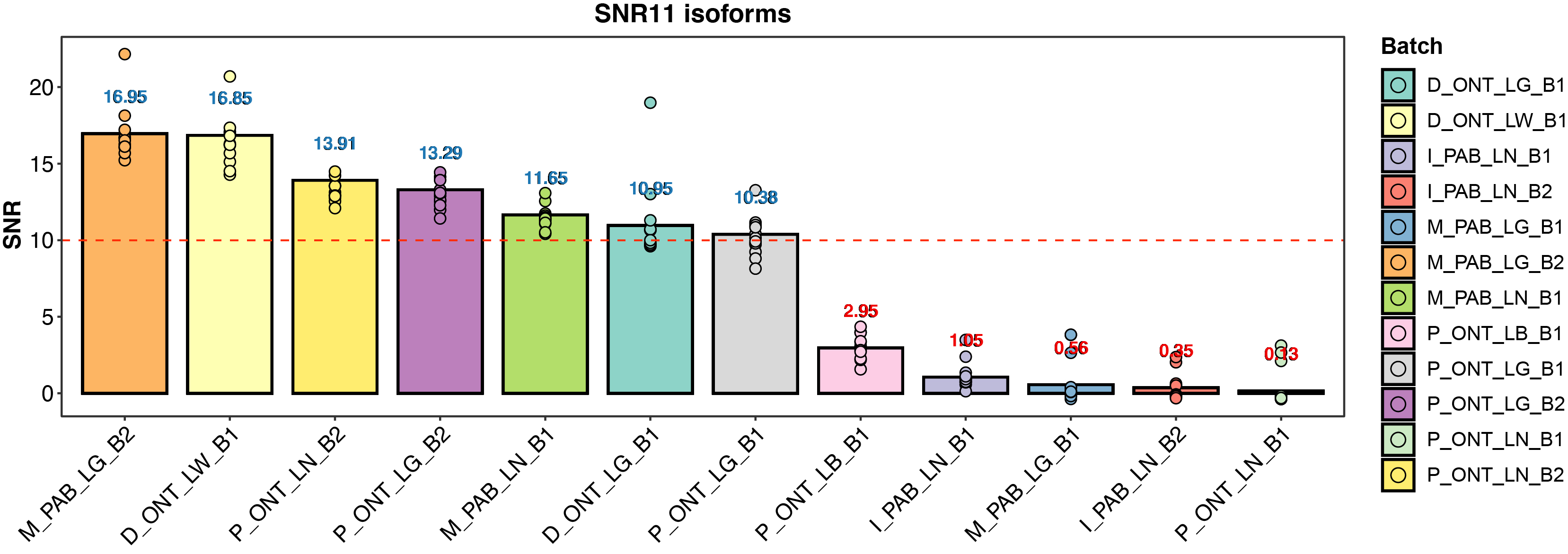

**Supplementary Fig. 13.** SNR across datasets based on the SNR and SNR11 criterion.

A barplot showing the SNR across datasets based on the SNR and SNR11 criterion for selecting high-quality datasets. Batches with SNR ≥ 10 meet the criterion and were selected for subsequent comparison with the srRNA-seq datasets.

### Supplementary Fig. 14. Plots with SNR values across 12 lrRNA-seq batches (Oarfish).

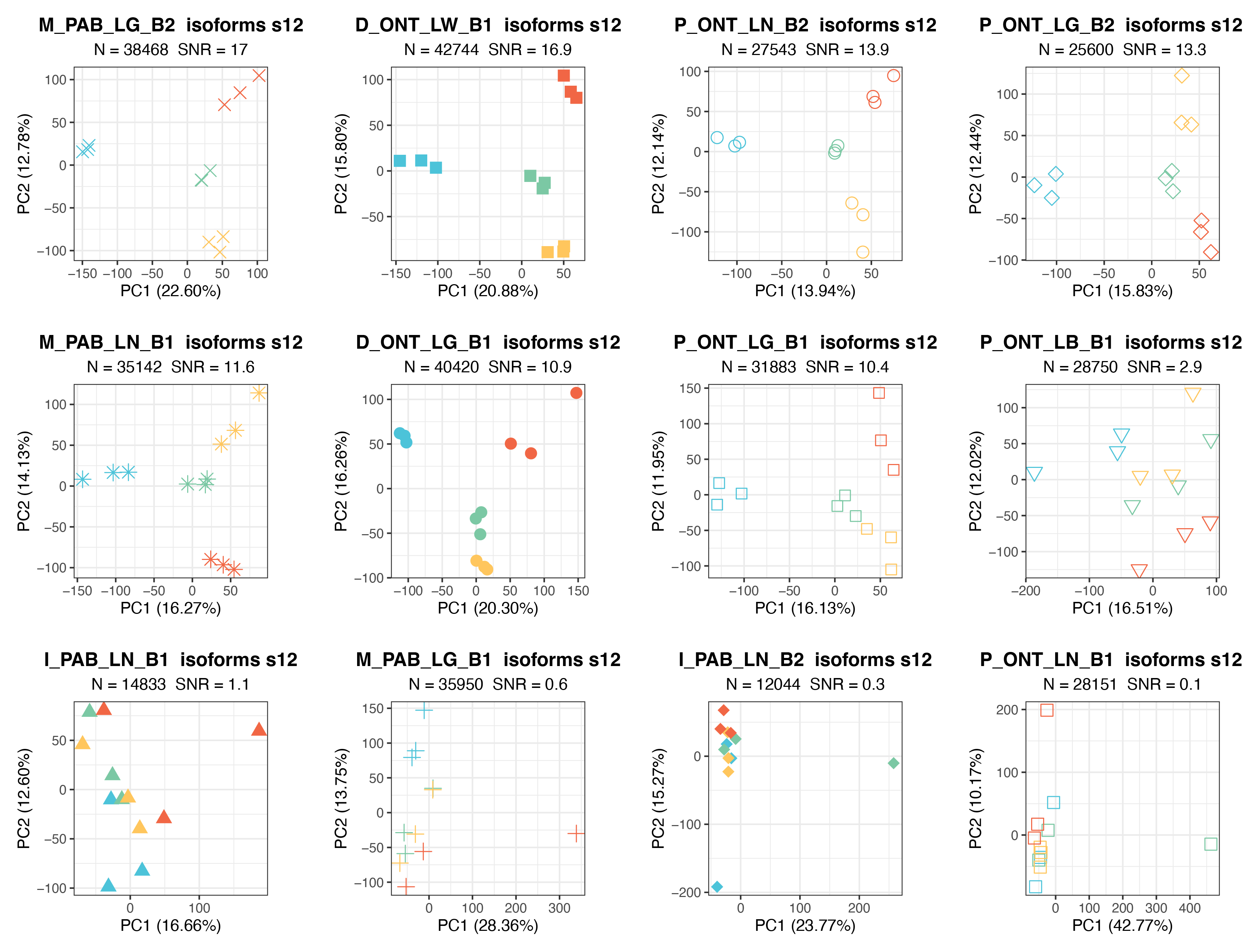

**Supplementary Fig. 14.** Plots with SNR values across 12 lrRNA-seq batches (Oarfish).

Each panel shows a PCA of absolute isoform abundances for one of the 12 batches (symbols denote batch; colors indicate sample type). The panel header reports the number of isoforms (N) retained after filtering and the corresponding SNR. Batches with SNR ≥ 10 meet the criterion and were selected for construction of the reference datasets, whereas the five low-SNR batches (bottom row) were excluded.

### Supplementary Fig. 15. SNR by isoform length for high-quality batches.

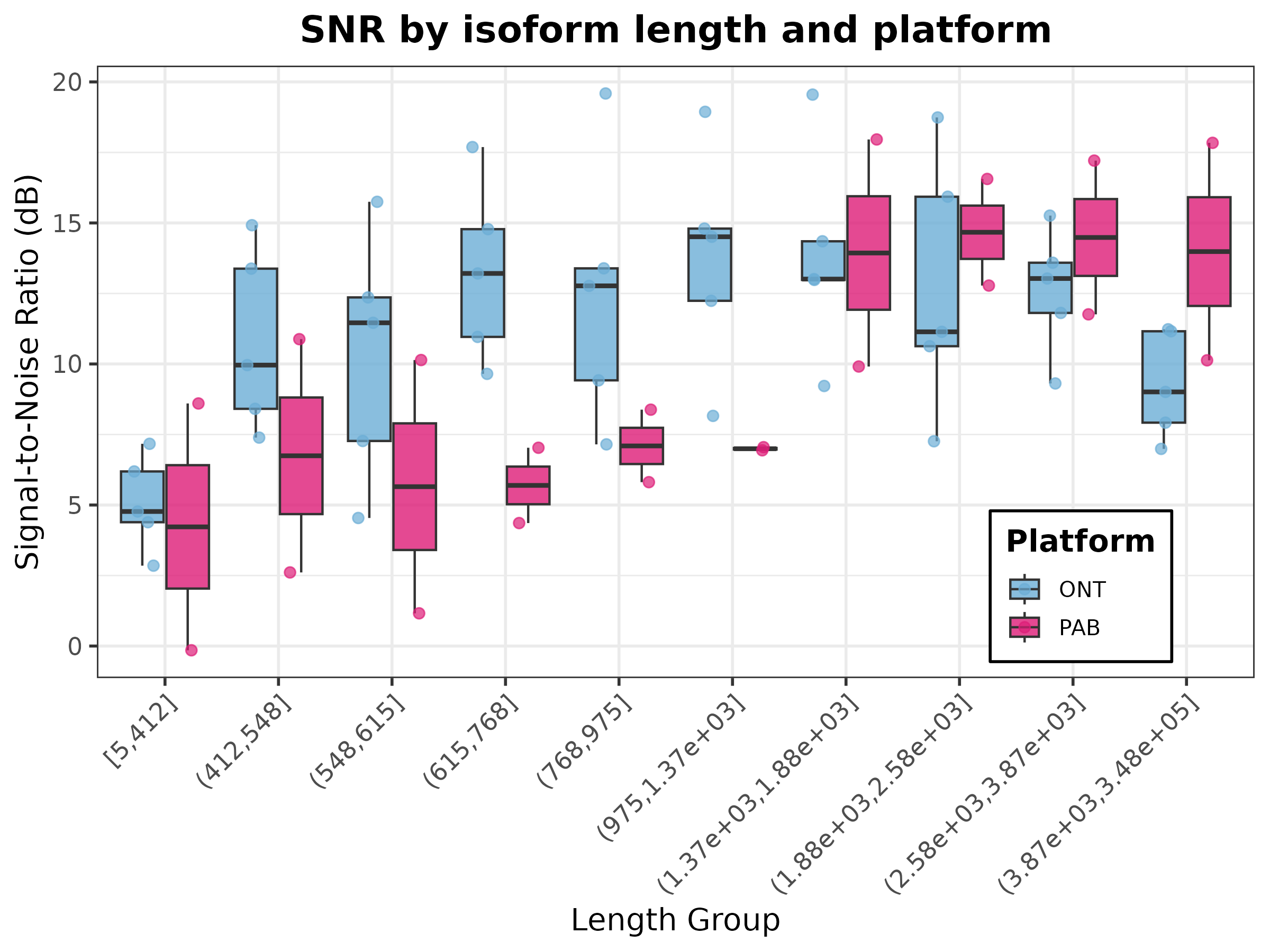

**Supplementary Fig. 15.** SNR by isoform length for high-quality batches.

Boxplots showing the SNR of absolute isoform expression for nine length bins (x-axis); bins are based on exon-spliced transcript length (bp). Data are derived from the seven high-quality batches retained. Colors distinguish sequencing platforms—Oxford Nanopore Technologies (ONT, blue) and PacBio (PacBio, magenta).

### Supplementary Fig. 16. Total reads across 22 short-read RNA-seq batches.

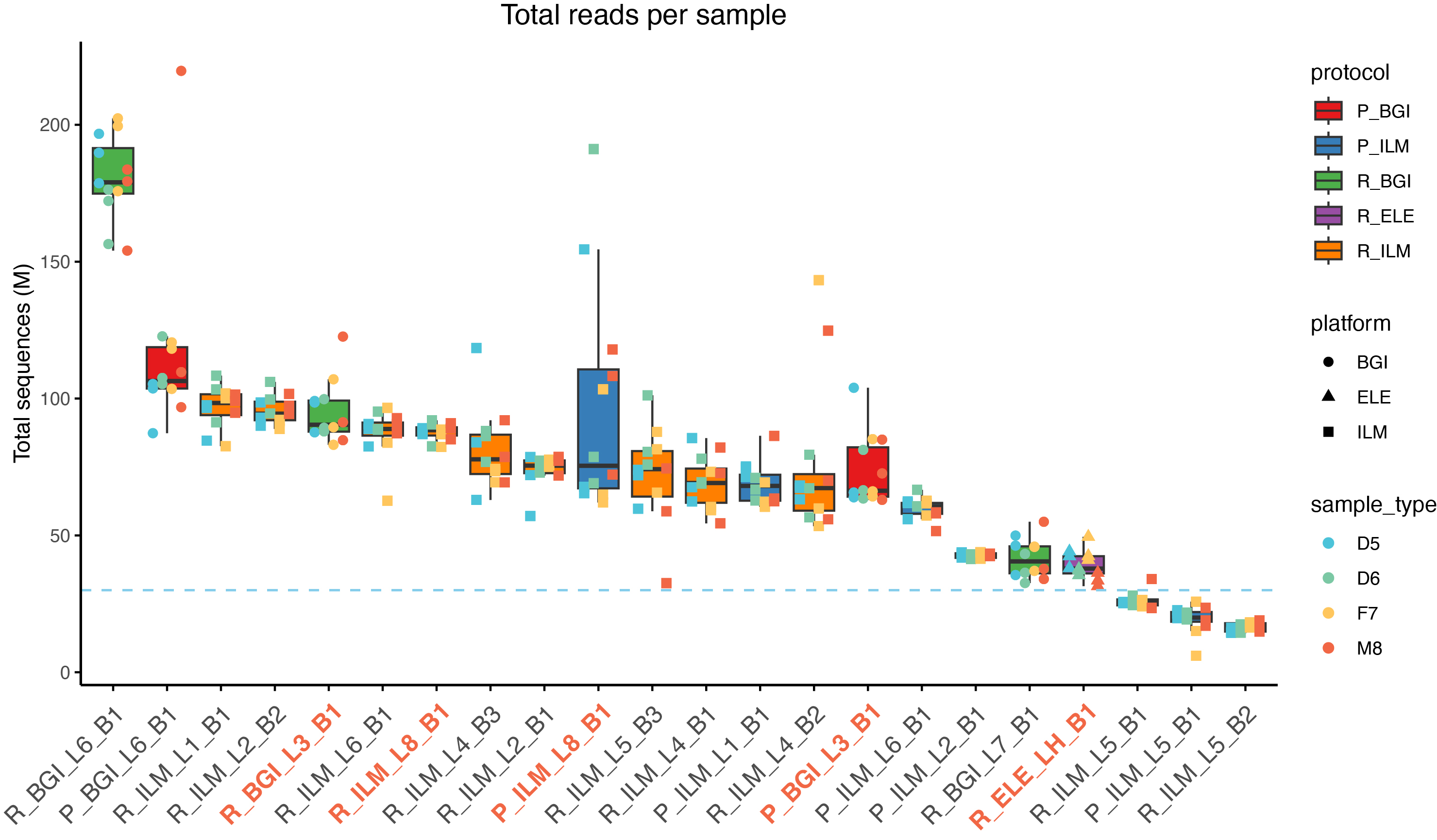

**Supplementary Fig. 16.** Total reads across 22 short-read RNA-seq batches.

Boxplots display the number of raw reads for every sample in each of the 22 short-read batches (y-axis in millions). Box fill colors distinguish library-prep protocols (P_BGI, P_ILM, R_BGI, R_ELE, R_ILM); point shapes mark sequencing platforms (circle, BGI; triangle, ELE; square, ILM); point colors indicate biological sample types (D5, D6, F7, M8). The dashed horizontal line marks the 30 M-read threshold adopted for downstream analyses. Batch names highlighted in red satisfy the coverage criterion and were retained for comparison with the lrRNA-seq datasets.

### Supplementary Fig. 17. Plots with SNR values across 22 srRNA-seq batches (Salmon).

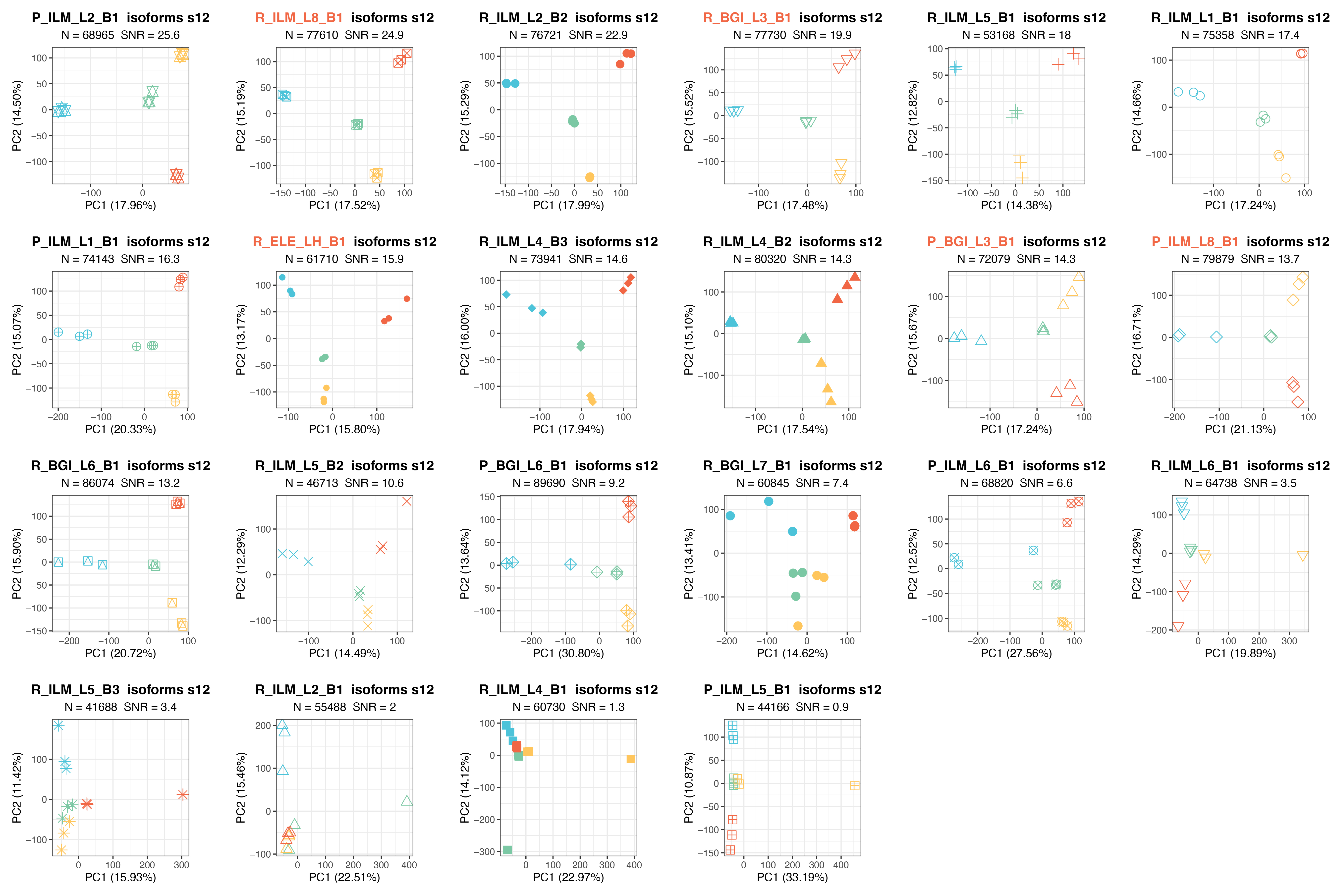

**Supplementary Fig. 17.** Plots with SNR values across 22 srRNA-seq batches (Salmon).

Each panel shows a PCA of absolute isoform abundances for one of the 22 batches (symbols denote batch; colors indicate sample type). The panel header reports the number of isoforms (N) retained after filtering and the corresponding SNR. Batches whose titles are highlighted in red were selected for direct comparison with the lrRNA-seq isoform datasets.

### Supplementary Fig. 18. Splice-junction complexity of the reference transcriptome.

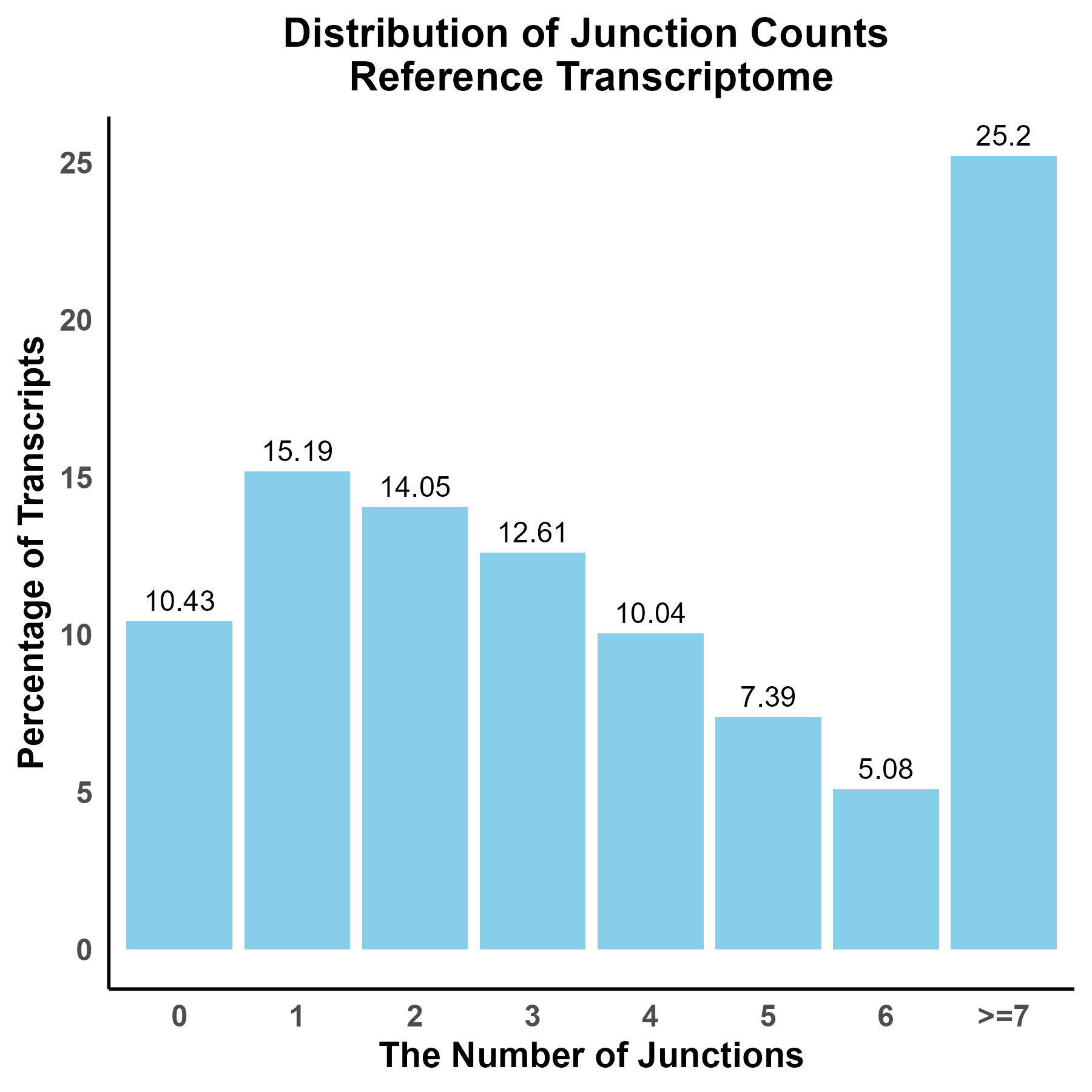

**Supplementary Fig. 18.** Splice-junction complexity of the reference transcriptome.

Histogram shows the percentage of reference transcripts that contain 0, 1, 2 … 6, or ≥ 7 intron–exon junctions. Single-exon (intron-less) transcripts account for 10%, whereas roughly one quarter of transcripts carry seven or more junctions, indicating that the reference set is enriched for multi-exon, structurally complex isoforms. Percentage values are printed above each bar for clarity.

### Supplementary Fig. 19. Global expression patterns before and after ratio normalization across lrRNA-seq and srRNA-seq.

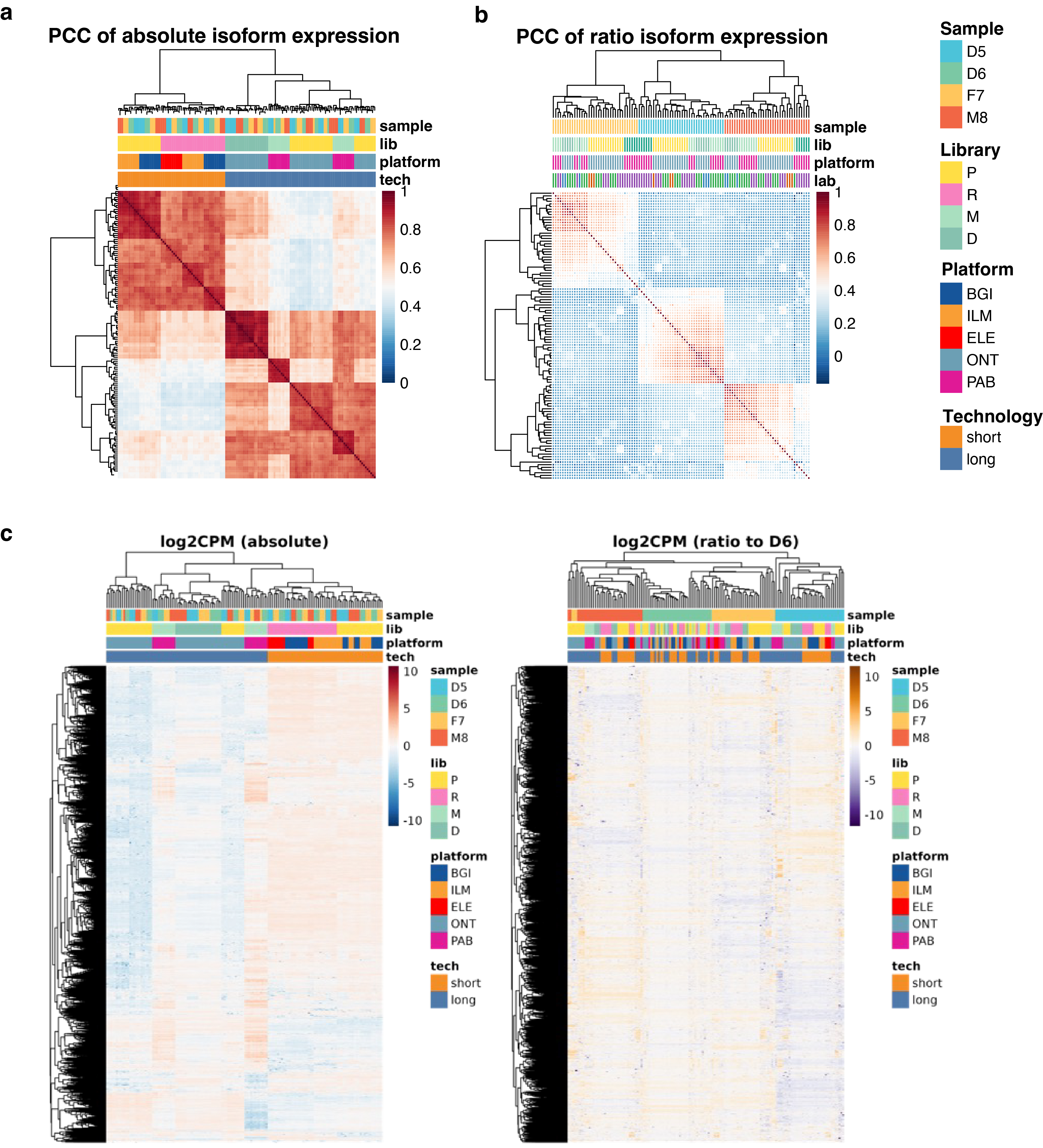

**Supplementary Fig. 19.** Global expression patterns before and after ratio normalization across lrRNA-seq and srRNA-seq.
**a**, Heatmap of pairwise Pearson correlation coefficients (PCC) for absolute isoform quantification across LR and SR datasets. Annotations indicate biological sample (color), library protocol, sequencing platform, and technology.

**b**, Heatmap of ratio-based isoform quantification, showing markedly higher cross-technology reproducibility.

**c**, Heatmaps display log₂-CPM values for the MANE isoforms across all long- and short-read libraries (rows). Columns are hierarchically clustered; colored bars annotate biological sample (D5, D6, F7, M8), library type (P, R, M, D), sequencing platform (BGI, ILM, ELE, ONT, PAB), and sequencing technology (short, long).

Left, absolute counts produce broad, protocol-driven blocks, with ONT and PacBio long-read datasets segregating from short reads.

Right, ratio normalization to the D6 replicate of each batch removes the platform effect: libraries now cluster primarily by biological sample regardless of technology, confirming effective cross-platform harmonization.

### Supplementary Fig. 20. PCA and PVCA analysis of four lrRNA-seq protocols and four analysis pipelines across 12 batches.

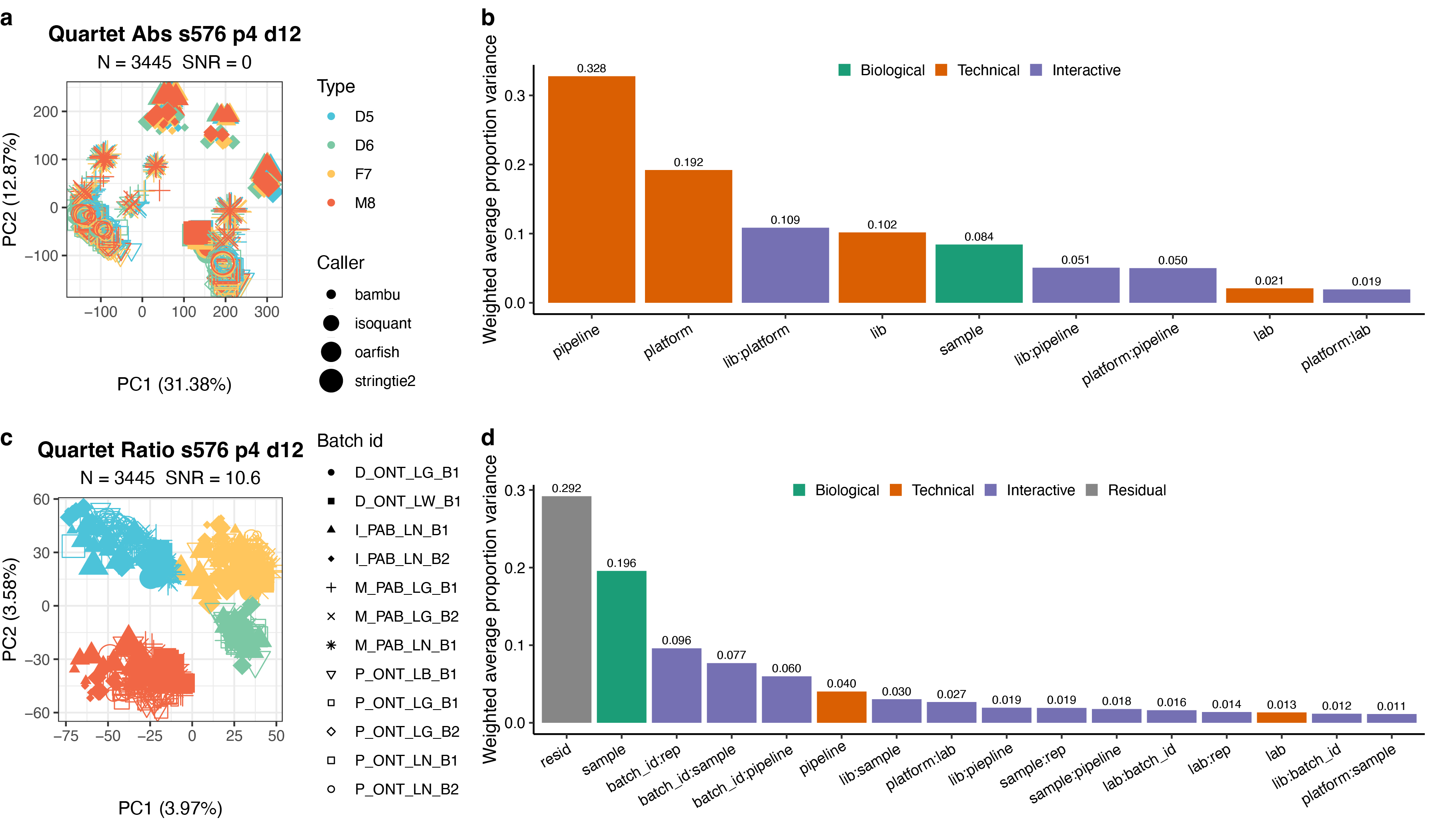

**Supplementary Fig. 20.** PCA and PVCA analysis of four lrRNA-seq protocols and four analysis pipelines across 12 batches.

**a**, Principal Component Analysis (PCA) plot using absolute expression values. Colors indicate sample type, sizes indicate the caller, and symbols denote the batch.

**b**, Principal Variance Component Analysis (PVCA) of absolute expression values showing variance proportions by technical (orange), biological (green), interactive (purple), and residual (grey) factors using absolute expression values.

**c**, Principal Component Analysis (PCA) plot using ratio-based expression values. Colors indicate sample type, sizes indicate caller, and symbols denote the batch.

**d**, Principal Variance Component Analysis (PVCA) of ratio-based expression values showing variance proportions.

### Supplementary Fig. 21. Reproducibility of direct-RNA m⁶A detection across batches.

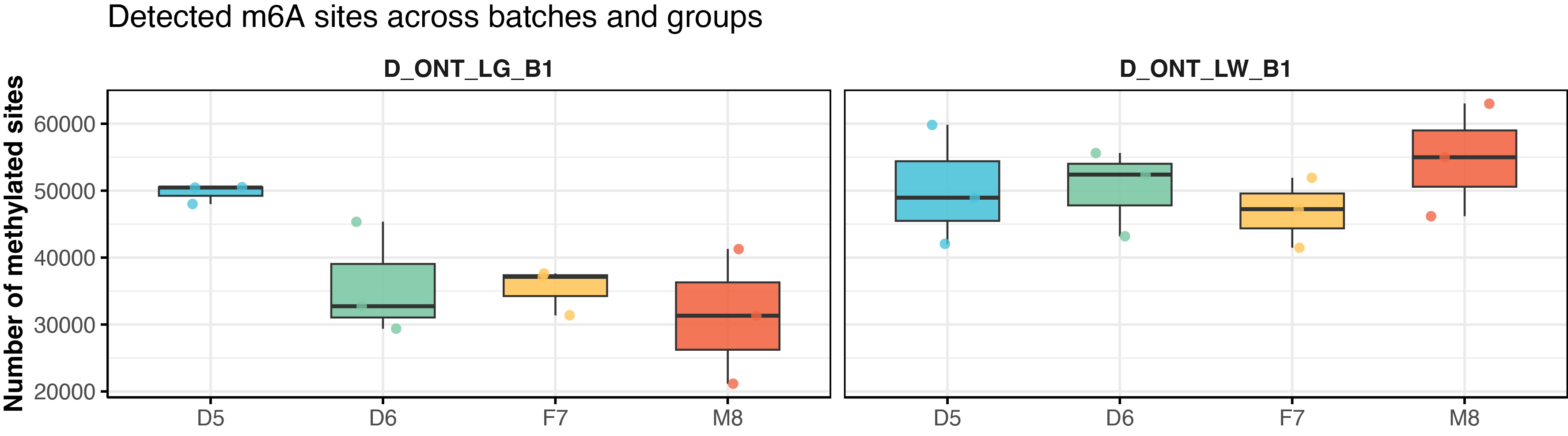

**Supplementary Fig. 21.** Reproducibility of direct-RNA m⁶A detection across batches.

Number of confidently called m⁶A sites per biological replicate (Supported modified counts ≥50). Boxplots are stratified by the two direct-RNA sequencing batches (D_ONT_LG_B1, left; D_ONT_LW_B1, right) and colored by sample (D5 cyan, D6 green, F7 yellow, M8 coral; center line = median, box = IQR, whiskers = 1.5 × IQR, points = outliers).

### Supplementary Fig. 22. Stepwise workflow for constructing benchmark reference datasets.

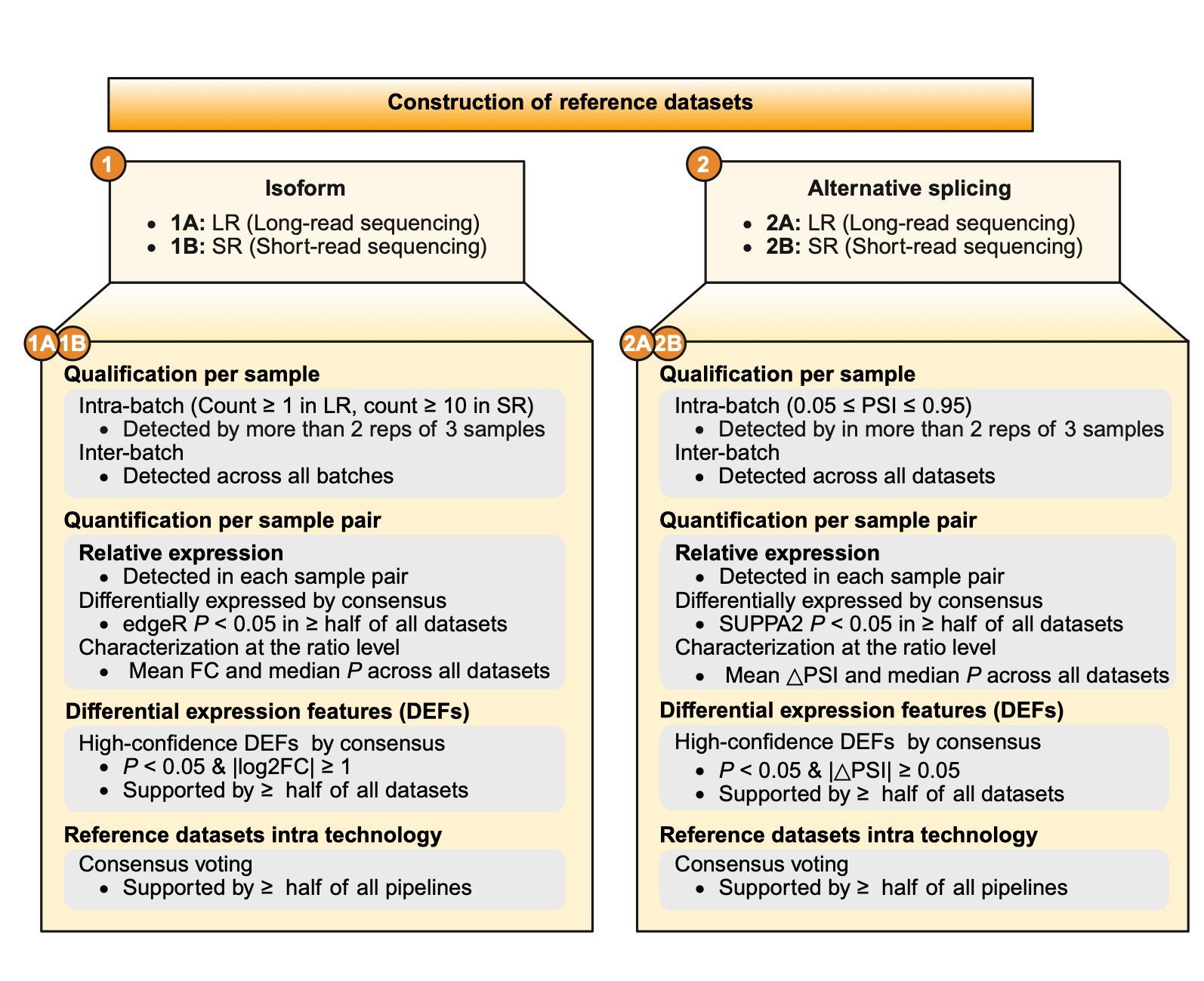

**Supplementary Fig. 22.** Stepwise workflow for constructing benchmark reference datasets.

The framework comprises two parallel branches: (1) isoform-level and (2) alternative-splicing (AS) level—each executed independently for long-read (LR) and short-read (SR) data (1A/1B and 2A/2B).

Qualification per sample. Isoforms/AS events are first filtered for robust detectability: they must be observed in ≥ 2 replicates within a batch and present in every batch of the same technology.

Quantification per sample pair. For every biological sample pair, we compute relative expression (ratio normalization). Differential events are called by consensus (edgeR for isoforms, SUPPA2 for AS) with *P* < 0.05 in ≥ 50% of datasets, and summarized by the across-dataset mean fold-change (FC) or ΔPSI.

High-confidence differential expression features (DEFs). Features meeting *P* < 0.05 and |log2FC| ≥ 1 (or |ΔPSI| ≥ 0.05 for AS) in ≥ 50% of datasets are retained as high-confidence DEFs.

Consensus reference sets. Finally, results from all analysis pipelines within the same technology are merged by majority vote; events supported by ≥ half of the pipelines constitute the LR and SR reference datasets used throughout the study.

### Supplementary Fig. 23. Schematic overview of qPCR validation.

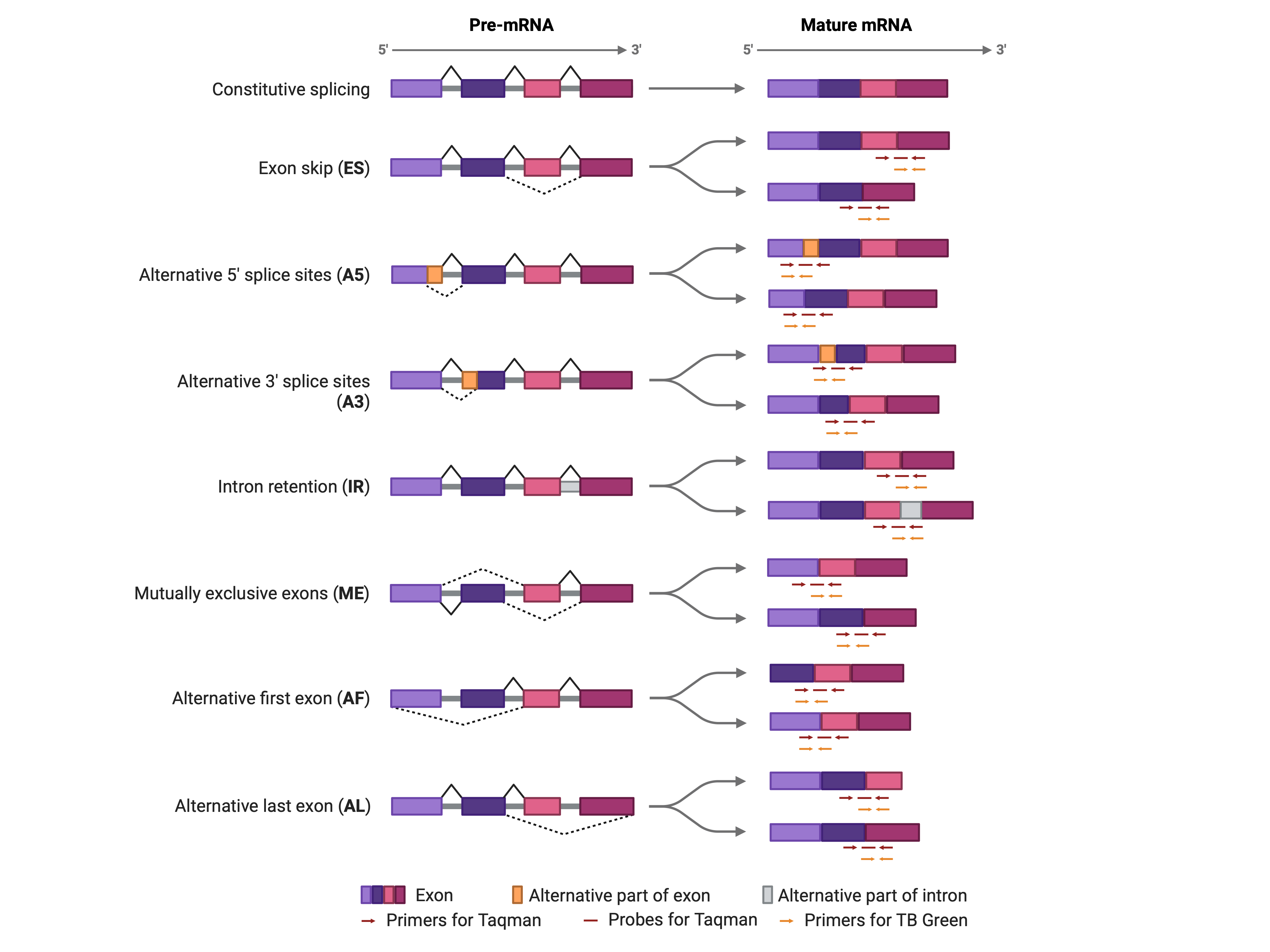

**Supplementary Fig. 23.** Schematic overview of qPCR validation.

To rigorously verify seven common alternative splicing events, this study employed RT-qPCR combining TB Green dye and TaqMan probe methods. Within a single alternative splicing event, a single gene generates two distinct isoforms, each containing a unique splicing junction. Primers and probes were designed to target these junctions, spanning the junction sites to enable relative expression quantification via Ct values. As illustrated, arrows and dashes below each isoform denote primer and probe target positions, serving as an intuitive reference for target sequence selection. Red arrows/dashes denote TaqMan primers/probes, while yellow arrows indicate TB Green primers.

### Supplementary Fig. 24. Concordance between long-only (LO) and short-only (SO) isoform reference datasets.

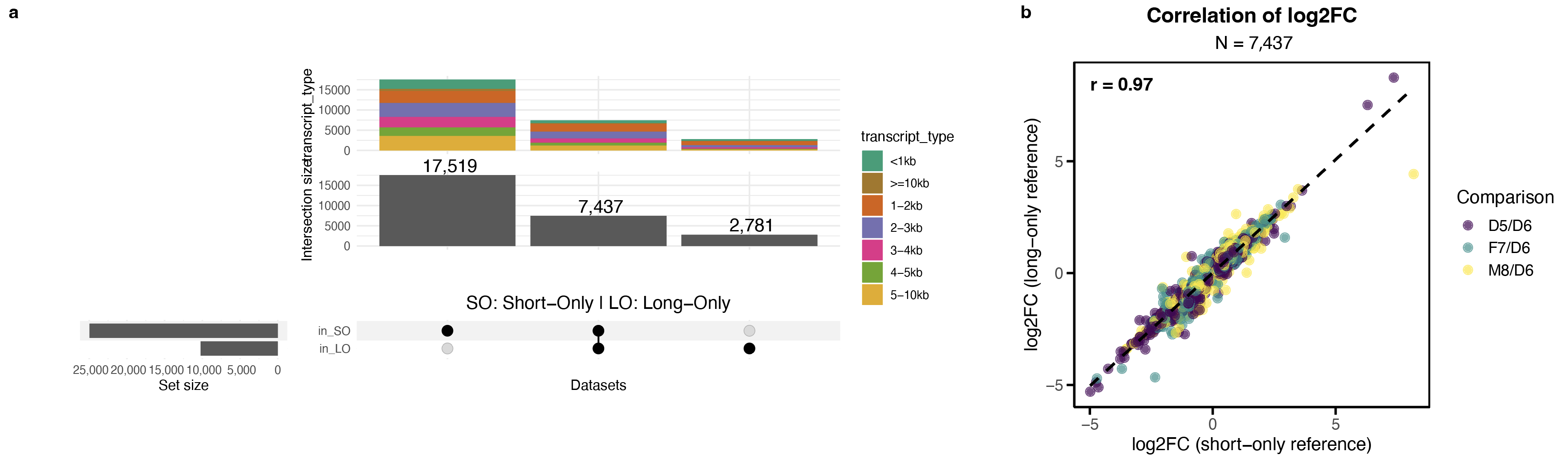

**Supplementary Fig. 24.** Concordance between long-only (LO) and short-only (SO) isoform reference datasets.

**a**, UpSet plot showing overlap between isoforms included in the SO and LO reference datasets. Among the total 27,737 isoforms, 7,437 were shared, 17,519 were SO-specific, and 2,781 were LO-specific. Bar colors indicate transcript length categories.

**b**, Correlation of log₂ fold changes (log₂FC) for shared isoforms (n = 7,437) between LO and SO reference datasets across three comparisons. A strong concordance is observed (Pearson’s *r* = 0.97), supporting the robustness of both datasets and the potential for integration.

### Supplementary Fig. 25. Concordance between long-only (LO) and short-only (SO) AS reference datasets.

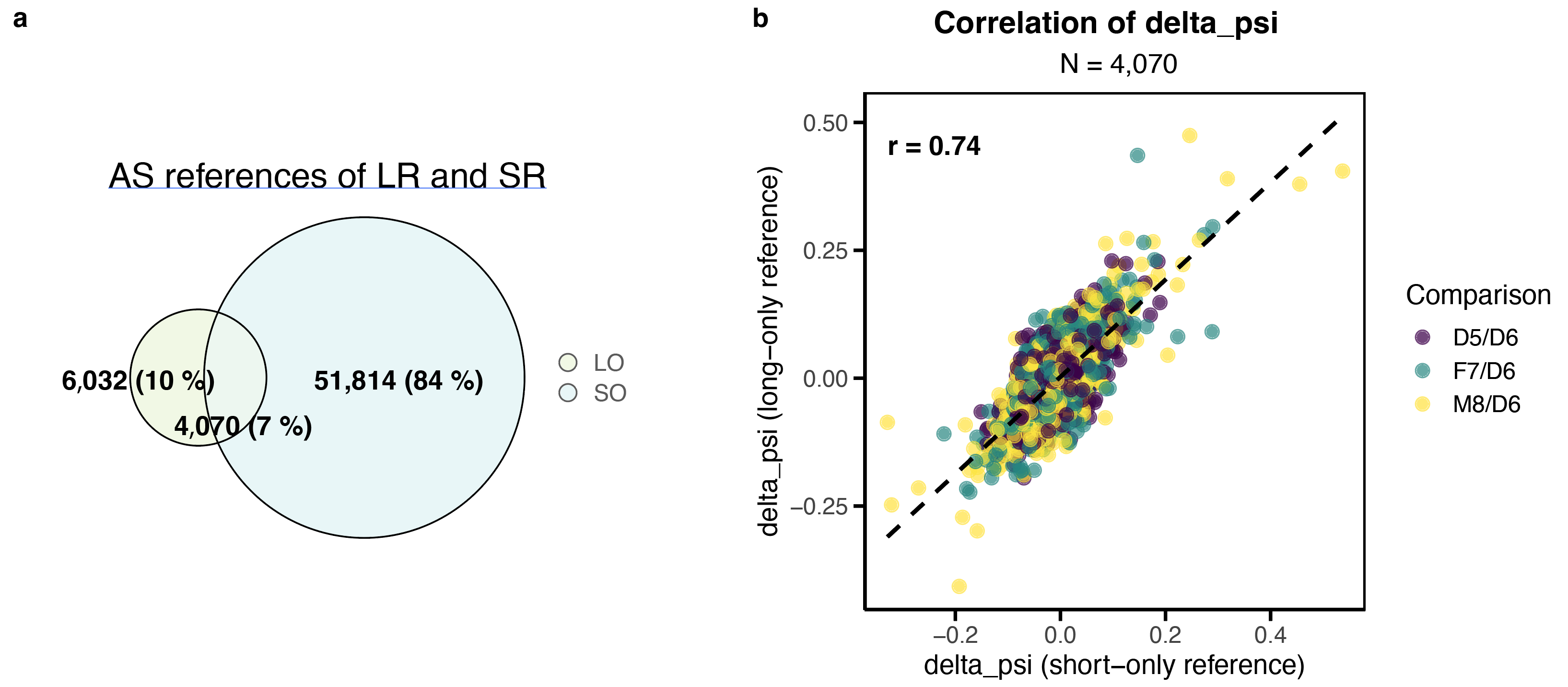

**Supplementary Fig. 25.** Concordance between long-only (LO) and short-only (SO) AS reference datasets.

**a**, Euler diagram showing the overlap of alternative splicing (AS) events identified in the LO and SO reference datasets. A total of 4,070 AS events (7%) were shared between LO and SO datasets, while 6,032 events (10%) and 51,814 events (84%) were identified exclusively in LO and SO datasets, respectively.

**b**, Scatter plot showing the correlation of ΔPSI values between LO and SO reference datasets for the 4,070 shared AS events across three comparisons (D5/D6, F7/D6, and M8/D6). Pearson correlation coefficient (*r* = 0.74) indicates high quantitative concordance between LO and SO reference datasets. The dashed line represents the fitted regression trend, illustrating the overall directional consistency between LO- and SO-derived ΔPSI values.

### Supplementary Fig. 26. Composite performance of quantification pipelines across the Quartet datasets.

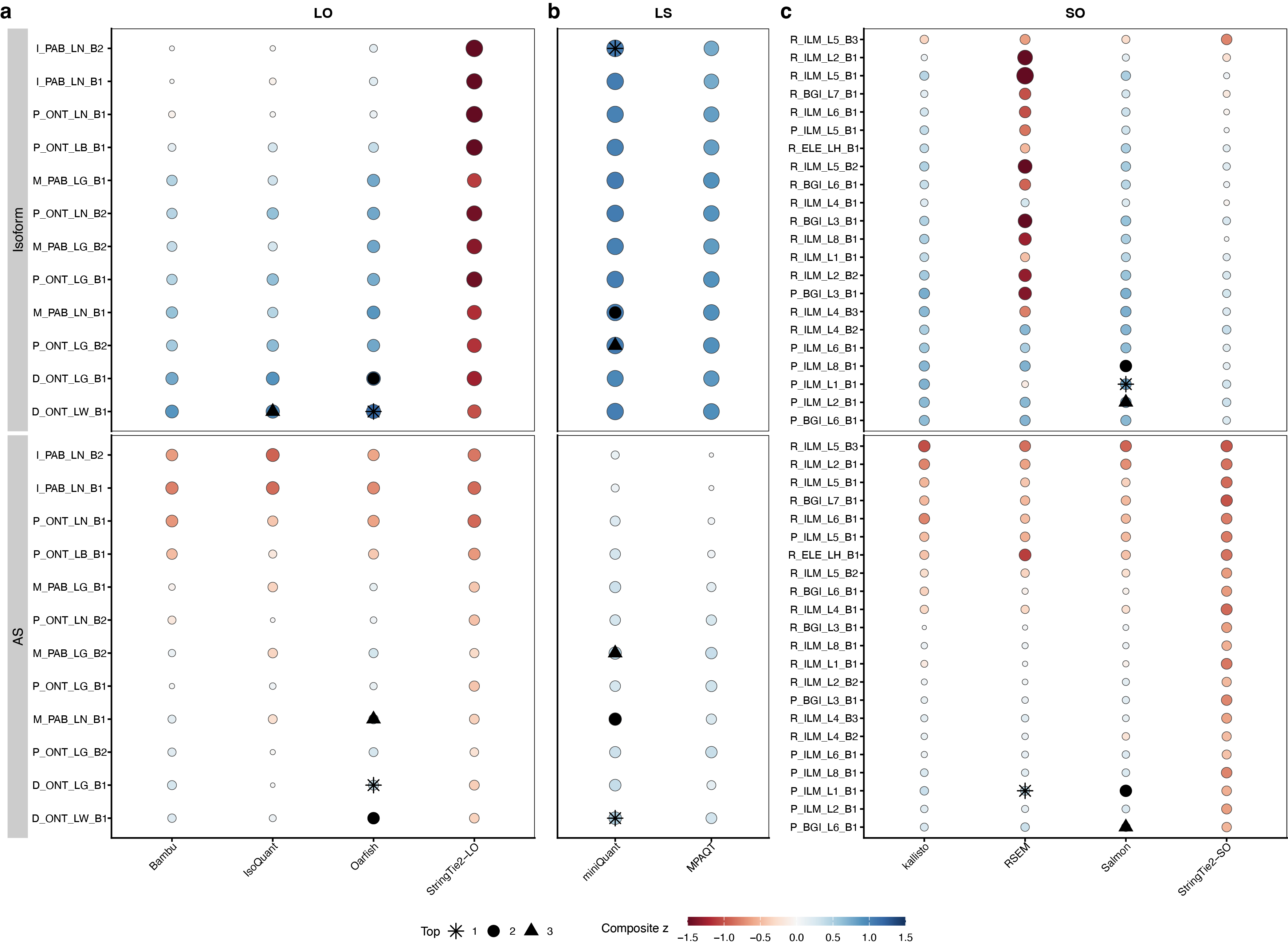

**Supplementary Fig. 26.** Composite performance of quantification pipelines across the Quartet datasets.

Bubble plots present the composite performance of **a**, long-read-only (LO); **b**, long + short hybrid (LS); and **c**, short-read-only (SO) pipelines. For each comparison group, the x-axis lists candidate pipelines, and the y-axis lists batches. Within every batch, a colored bubble is plotted separately for isoform (upper facet) and AS (lower facet) quantification. The bubble fill encodes the composite z-score that averages three accuracy metrics: Pearson correlation (RC), root mean square error (RMSE; sign-reversed), and Matthews correlation coefficient (MCC). Blue indicates above-average performance, red below-average, and white the global mean. Bubble size is proportional to |z|, emphasizing stronger departures from the mean. Stars, circles, and triangles label the first-, second-, and third-ranked pipelines within each facet and compare-group; pipelines lacking results for a given batch are not shown.

### Supplementary Fig. 27. Assembly of the Quartet-specific transcriptome.

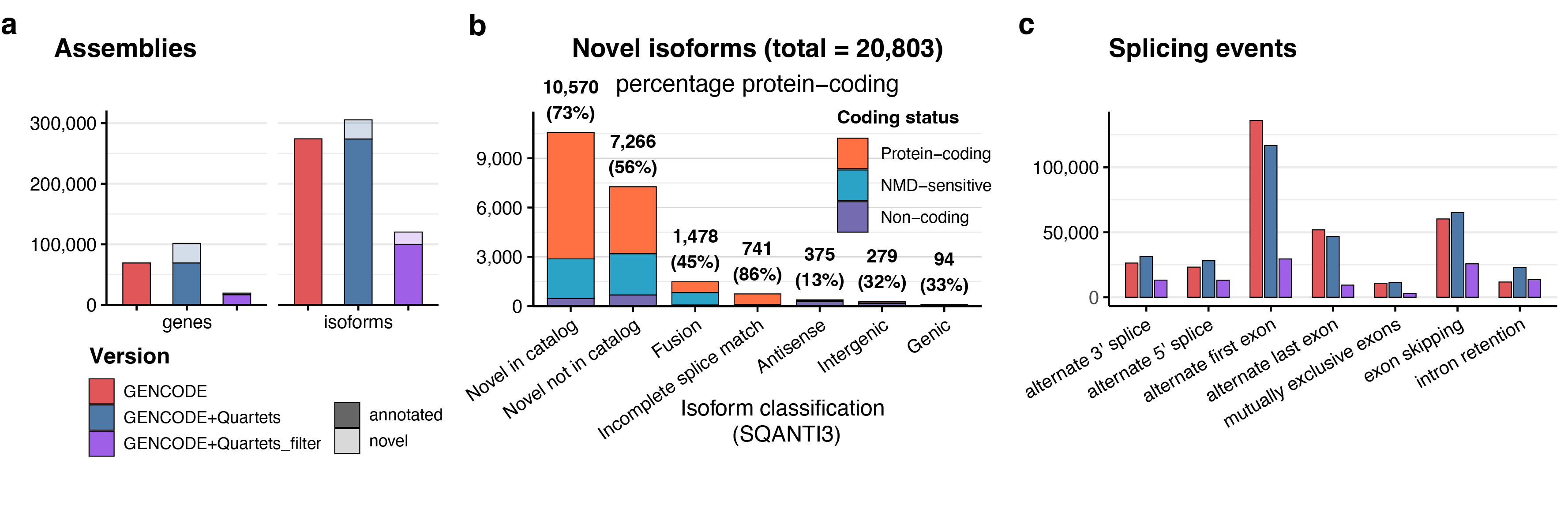

**Supplementary Fig. 27.** Assembly of the Quartet-specific transcriptome.

**a**, Growth of the Quartet assembly when GENCODE is augmented with long-read data (GENCODE + Quartets) and after high-confidence filtering (GENCODE + Quartets_filter); bars are split into annotated (dark) and novel (light) isoforms.

**b**, SQANTI3 structural classification of the 20,803 novel isoforms. Stacked colors indicate predicted coding status (protein-coding, NMD-sensitive, non-coding), and percentages above bars give the fraction protein-coding.

**c**, Catalogue of alternative-splicing events in the three assemblies, showing the marked contribution of the hybrid (GENCODE + Quartets_filter) set to retained-intron and mutually-exclusive exon annotations.

### Supplementary Fig. 28. Comparison of transcriptome assemblies between StringTie3 and Bambu.

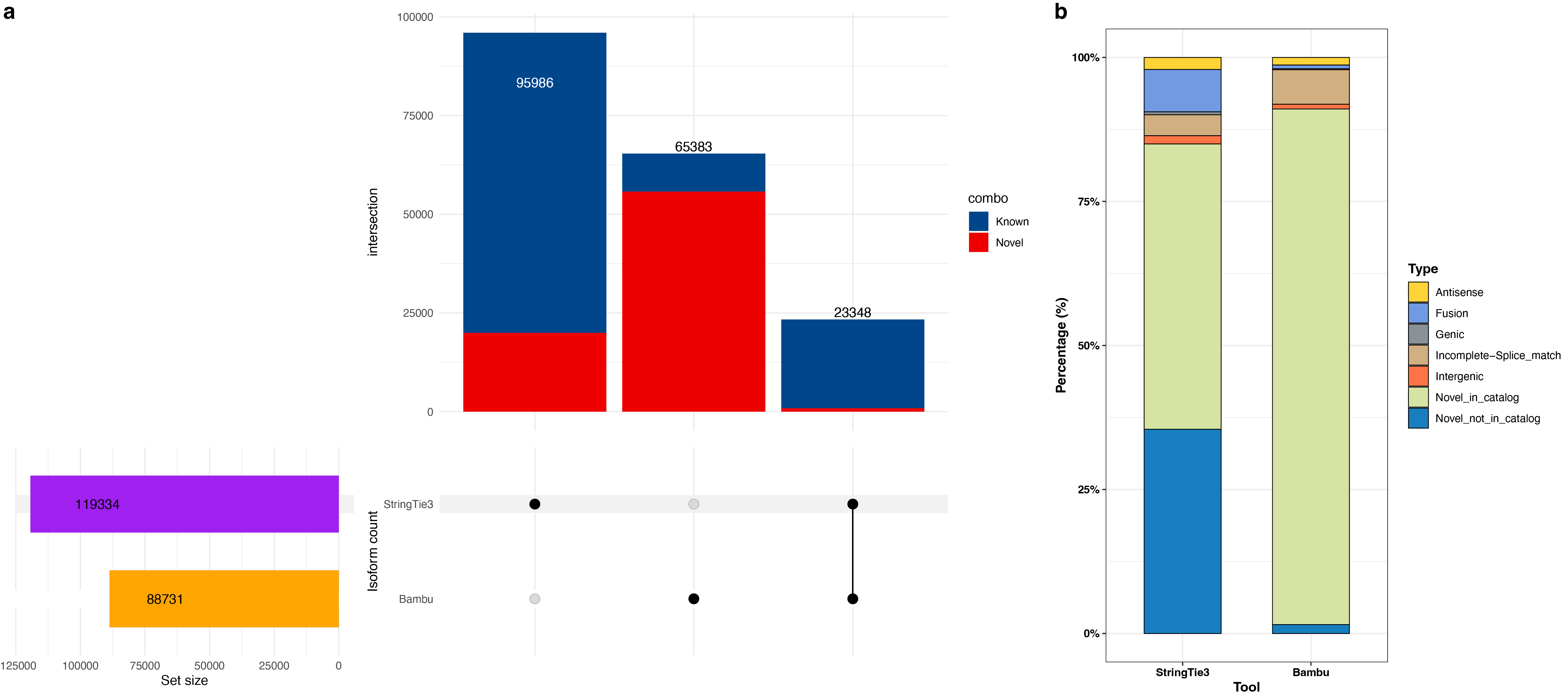

**Supplementary Fig. 28.** Comparison of transcriptome assemblies between StringTie3 and Bambu.

**a**, UpSet plot showing the overlap between the two assemblies. Horizontal bars on the left give the total number of transcript models output by each tool (StringTie3, 119,334; Bambu, 88,731). Vertical bars above the matrix indicate the number of isoforms found only by StringTie3 (95,988), only by Bambu (22,438), or by both tools (65,343). Bar fill distinguishes GENCODE-annotated (‘known’, blue) from novel (‘new’, red) isoforms, showing that most tool-specific models are novel, whereas the shared set is enriched for annotated transcripts.

**b**, SQANTI3 classification of novel isoforms recovered by each assembler. Both tools are dominated by novel-in-catalog (NIC) and novel-not-in-catalog (NNC) transcripts, with smaller but comparable fractions of antisense, intergenic, genic, fusion, and incomplete-splice-match (ISM) classes.

### **Supplementary Tables**

### Table S1. Metadata of the Quartet lrRNA-seq datasets.

### Table S2. Benchmark source comparison.

### Table S3. Metadata of the Quartet srRNA-seq datasets.

### Table S4. Performance metrics overview.

### Table S5. Ratio-based isoform reference datasets.

### Table S6. Ratio-based differentially expressed isoforms (DEIs) reference datasets.

### Table S7. The log2 fold changes and *p*-values of reference isoforms tested in reference datasets and qPCR results.

### Table S8. Ratio-based alternative splicing (AS) reference datasets.

### Table S9. Ratio-based differential alternative splicing events (DASEs) reference datasets.

### Table S10. The mean delta psi and *p-*values of reference isoforms tested in reference datasets and qPCR results.

### Table S11. Primers for validated isoforms for TB Green method.

### Table S12. Primers and probes for validated isoforms for TaqMan method.

### Table S13. Primers for validated alternative splicing events (ASEs) for TB Green method.

### Table S14. Primers and probes for validated alternative splicing events (ASEs) for TaqMan method.
